# A spatiotemporal gradient of mesoderm assembly governs cell fate and morphogenesis of the early mammalian heart

**DOI:** 10.1101/2022.08.01.502159

**Authors:** Martin H. Dominguez, Alexis Leigh Krup, Jonathon M. Muncie, Benoit G. Bruneau

**Affiliations:** Gladstone Institutes, San Francisco, CA, USA; Department of Medicine, Division of Cardiology, University of California, San Francisco, CA, USA; Cardiovascular Institute, University of Pennsylvania, Philadelphia, PA, USA; Biomedical Sciences Graduate Program, University of California, San Francisco, CA, USA; Roddenberry Center for Stem Cell Biology and Medicine at Gladstone, San Francisco, CA, USA; Department of Pediatrics and Cardiovascular Research Institute, University of California, San Francisco, San Francisco, CA, USA

## Abstract

Using four-dimensional whole-embryo light sheet imaging with improved and accessible computational tools, we longitudinally reconstruct early murine cardiac development at single-cell resolution. Nascent mesoderm progenitors form opposing density and motility gradients, converting the temporal birth sequence of gastrulation into a spatial anterolateral-to-posteromedial arrangement. Migrating precardiac mesoderm doesn’t strictly preserve cellular neighbor relationships; spatial patterns only become solidified as the cardiac crescent emerges. Progenitors undergo a heretofore unknown mesenchymal-to-epithelial transition, with a first heart field (FHF) ridge apposing a motile juxtacardiac field (JCF). Anchored along the ridge, the FHF epithelium rotates the JCF forward due to push-pull morphodynamics of the second heart field, which forms the nascent heart tube. In *Mesp1* mutants that fail to make a cardiac crescent, mesoderm remains highly motile but directionally incoherent, resulting in density gradient inversion. Our practicable live embryo imaging approach defines spatial origins and behaviors of cardiac progenitors, and identifies their unanticipated morphological transitions.

## Introduction

The emergence and allocation of the progenitors of organs offers insights into the events that ensure robust morphogenesis. The developing heart is particularly sensitive to disturbed morphogenesis, as congenital heart defects occur in over 1% of live births. Understanding the stepwise allocation and assembly of cardiac precursors will provide insights into heart development and disease. Cell labeling and histological studies have shown how the heart forms from its earliest discernible stages [1–5], but individual cellular events following gastrulation remain mostly uncharacterized.

Cardiovascular progenitors emerge during gastrulation as a subset of the *Mesp1*^+^ nascent mesoderm population, and migrate to lateral regions that will become the cardiac crescent [6–8]. Early cardiac progenitors comprise multipotent progenitor pools, the first and second heart fields (FHF and SHF), as well as a newly-classified juxta-cardiac field (JCF). The JCF contributes to epicardium and left ventricle (LV) [9,10]. Partially overlapping the JCF, the FHF contributes to atria, atroventricular canal (AVC) and left ventricle (LV) [6,11]. SHF cells contribute to the atria, right ventricle (RV), and outflow tract (OFT) [12,13].

Mouse genetics tools have led to complex lineage and clone labeling strategies, revealing that *Mesp1*^+^ progenitors have rudimentary assignments to final cardiac structures, even prior to formation of the heart fields. Notably, both temporal and spatial restriction of the *Mesp1*^+^ progenitor pool have been shown [6,7,14]. However, evidence that may unify our understanding of early specification in both temporal and spatial domains, is incomplete. Moreover, due to the complex morphological processes that underpin heart formation, concretely linking early progenitors to their progeny structures requires examination at greater temporal resolution than lineage tracing alone can afford.

Live imaging of avian cardiogenesis has brought insights in early cardiac morphogenesis, exploiting the relative accessibility of such embryos for visualization and micro-manipulation [2,15–17]. Imaging studies of early mouse development, however, have grown at a relatively slower pace, owning to the fragility and limited longevity of *ex vivo* embryo culture [18–21]. Recent studies have examined gastrulation [22] and cardiogenesis [23] in the mouse, but are limited to examining only a few cells at a time.

Light sheet fluorescence microscopy (LSFM) is well suited to morphogenetic studies of mouse development [24–27], though most in toto embryo imaging has been performed on highly-specialized, custom-build instruments. While computational analysis of large-scale LSFM data is now possible [27– 30], most existing software applications are designed with the same specialization as the custom microscopes with which they are paired.

Overcoming these roadblocks, we performed comprehensive whole-embryo analyses to examine early cardiac progenitors and their emergence from *Mesp1*^+^ mesoderm. We combined a widely-available LSFM setup and murine *ex vivo* embryo culture (Fig. 1A), integrating data from fluorescent reporters for both *Mesp1* lineage and the *Smarcd3 “*F6” enhancer, the latter being the earliest known cardiac-specific identifier [6]. Furthermore, we generated new computational tools and improved existing ones, aiming to enhance data collection, image processing, and computational analysis of such large-scale data, and to help democratize the use of live embryo imaging.

**Figure 1:**
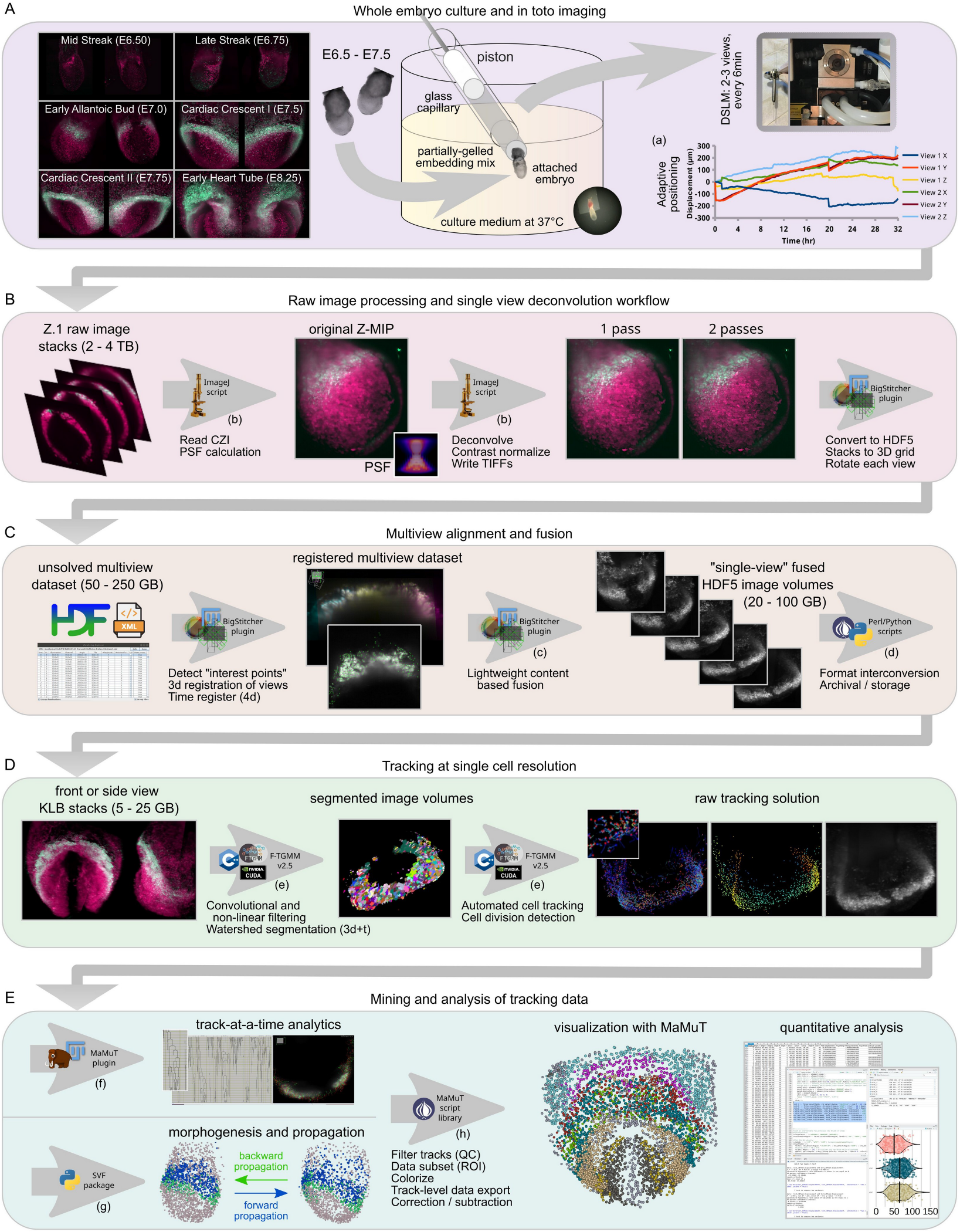
Comprehensive workflow for quantitative analysis of embryogenesis by live LSFM. **A**. Biology and microscopy protocol. Red signal: *Mesp1* lineage, green signal: *Smarcd3*-F6-nGFP. ZLAPS adaptive positioning example is shown, demonstrating shifts of two views’ XYZ positions needed to maintain the embryo in the center of the view. **B**. Initial raw .czi computational workflow, depicting macro-based batch deconvolution, filtering, and image import to BigStitcher. **C**. Multiview alignment to generate single image volume for each channel and timepoint. **D**. Improved tracking with F-TGMM v2.5. **E**. F-TGMM results can be refined using SVF, generating long-track morphodynamic models, or can be examined raw. MaMuT Perl script library annotates, filters, subsets, combines, and exports data. Lower case letters correspond with repositories listed in Software table within materials and methods.

By tracking cardiogenesis at single cell resolution with retrospective *in silico* labeling, our work reveals how cardiac regional fate is intimately tied to the temporal birth and migration sequence of cardiac progenitors. Additionally, we highlight the morphological formation of cardiac epithelium, uncovering region-specific migration and movement behaviors that ultimately shape and sculpt the early heart.

## Results

### An improved computational workflow for in toto mouse embryogenesis by multi-view LSFM

Cardiac fate allocation occurs early in gastrulating embryos [6,7]. We explored means of live investigation using fluorescent reporter mice that may uncover mechanisms underlying the genesis of early cardiac progenitors and their allocation to the heart tube.

Recently, McDole et al. described a comprehensive, whole-embryo imaging workflow of mouse post-implantation development [29]. The powerful LSFM microscope utilized in that study provides unparalleled imaging, but assembly time can range from weeks to months, the instrument occupies an entire room, and it requires dedicated specialists to operate. As an alternative, cost-effective but advanced commercial LSFM setups such as the Zeiss Lightsheet Z.1 are becoming widely available. To facilitate long imaging runs on our Z.1, we wrote an interactive application to perform adaptive position correction by registering sequential captures and interfacing with the microscope’s software (Fig. 1A).

We empirically determined that 2-3 specimen views acquired at 6-minute intervals would produce an acceptable balance of data return and phototoxicity, and sought a compatible computational pipeline for downstream analysis. Raw data amounts to 2-4 terabytes per experiment, depending on number of views, channels, and duration. A true in toto approach then “fuses” those views to form a single comprehensive image volume of the entire specimen, with deblurring methods applied in the process [31].

One such method, multiview deconvolution, becomes computationally efficient with 4 or more views [32]. As we utilize only 2-3 oblique views of each embryo, we crafted an open-source single-view deconvolution and fusion workflow (Fig. 1B), avoiding iterative methods due to their staggering processing overhead with this type of data. Our macro-based application employs closed-form deconvolution [33] in batch (using theoretical PSFs), offering further enhancement with Fiji’s background subtraction algorithm [34]. We next employed BigStitcher [28], a user-friendly tool for registering (i.e. aligning) and fusing (consolidating) multiview LSFM datasets in 4d (Fig. 1C).

Within BigStitcher, we carefully examined content-based fusion, a method that vastly outperforms mean fusion in terms of result quality (Fig. S1A). It does this by estimating regions of entropy (i.e. noise) in each view, and weighting the output to favor entropy-low regions. However, content-based fusion is impractical or even unattainable with large datasets due to its memory and CPU consumption. Since the weight images for each view are, in effect, compacted summaries of the content within the image, we reasoned that downscaling prior to entropy calculation may have little effect on either the weight images or the fusion results. Indeed, 2X or 4X downscaling (prior to entropy calculation) produced nearly identical results across a wide range of sample datasets, but with markedly decreased CPU time and memory usage (Fig. S1A’-A”). We named the optimized algorithm “lightweight” content-based fusion (Fig. 1C).

After multiple views are consolidated into a single volume for each channel and timepoint, tracking is used to estimate each cell’s position in space and time. We started with open source TGMM 2.0 [29], adding several enhancements to tracking accuracy and computational efficiency (Fig. S1B). We first improved TGMM’s segmentation by employing a dynamic background subtraction routine, utilizing image features (derived from Gaussian blur filtering) to identify background, rather than by subtracting static pixel values homogeneously (Fig. S1C-D’). Next, we optimized the main tracking loop to minimize repeat calls to hierarchical segmentation by caching their results. Finally, we re-wrote the division detection machine learning classifier to score combinatorial division trios (mother-daughter-daughter) near each cell birth, choosing the best trio for the final solution (Fig. S1H-I). With its ultimate iteration designated v2.5, Forked Tracking with Gaussian Mixture Models (F-TGMM) represents a stabler and more accurate tracking package (Fig. S1G) that runs 30% faster than TGMM 2.0 (Fig. S1B’).

TGMM data can be analyzed as raw tracks (Fig. 1E), which spuriously and stochastically terminate owing to imperfect linkage across time (Fig. S1E-G). Alternatively, tracks can be extended in time to create a morphodynamic overview of the dataset, using a package called statistical vector flow (SVF, Fig. 1E) [29]. The open source Fiji plugin, MaMuT, is used for visualization of raw tracks and SVF results [30]. We updated SVF for use with Python 3 (Python 2.x is no longer maintained), and enhanced MaMuT for 3d viewing of large datasets, and for displaying cell vector flow (Fig. 1E). Lastly, we wrote a collection of scripts for MaMuT dataset manipulation (Fig. 1E), which perform a variety of tasks that: selectively label or color embryo regions/tissues, subset and concatenate datasets, export track data for statistical analysis, filter or exclude tracks by cell or track features, and more.

Overall, these computational tools facilitate collection, analysis, and visualization of in toto live imaging data. We applied this comprehensive package to the investigation of mesoderm migration and early cardiac morphogenesis, though it could be used in a variety of applications. All are open source and portable, and compatible with contemporary hardware and software.

### A spatiotemporal gradient of mesoderm accumulation

After finalizing the computational toolbox for live imaging of mouse embryos, we examined behavior during and immediately following gastrulation as cardiac progenitors are born. Using *Mesp1* lineage reporter mice, we began in toto experiments at the mid streak (E6.5) stage, when only a few progenitors have arrived in the mesoderm layer (Fig. 2A and Video S1). Across all embryos studied, we noticed stereotypical collective migration of the mesoderm, yet stochastic individual cell behaviors.

**Figure 2:**
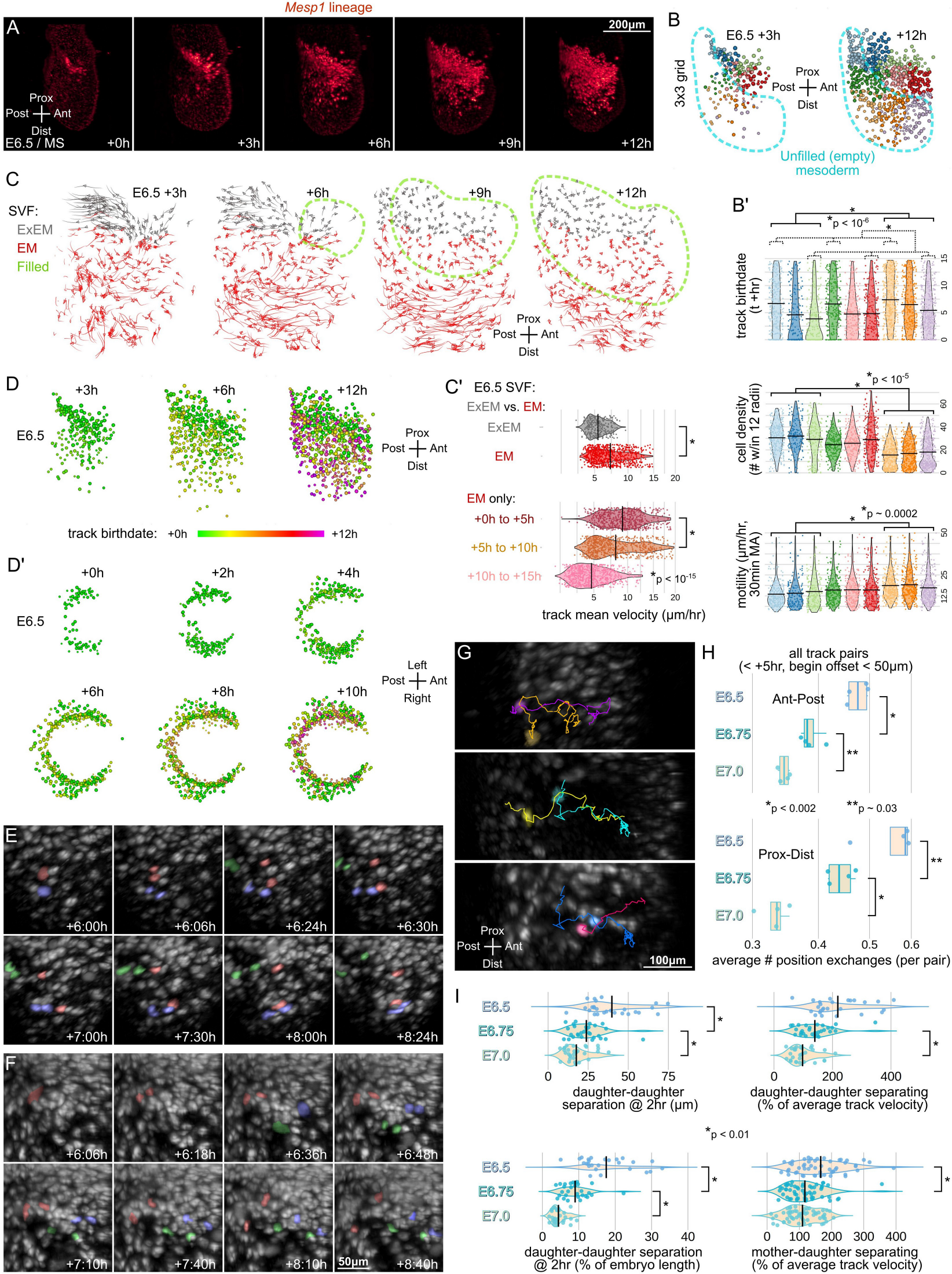
A spatiotemporal gradient of mesoderm accumulation. **A**. Time-lapse whole-embryo LSFM imaging of *Mesp1* lineage at E6.5, showing lateral view right-half max projections. **B**. Side views of a TGMM/MaMuT reconstructed E6.5 embryo during live imaging, with all tracks retrospectively partitioned onto a 3×3 grid. **B’**. TGMM tracks analyzed from +0h to +15h for birthdate, motility, and cell density. **C**. TGMM/SVF reconstruction of E6.5 anterior mesoderm migration, in orthographic projection with uniform sparsification. Extraembryonic (ExEM) and embryonic (EM) compartments are colored. **C’**. Quantification of this SVF series. **D-D’**. TGMM reconstruction E6.5 embryo live imaging, cells painted by track birthdate. **E-F**. Three manually annotated, randomly dispersed, division events are shown in each **E** and **F**, overlayed in false color (division nodes and daughter cells) on single sided, lateral Z-projections from live imaging experiments at E6.5. **G**-**H**. Analysis of TGMM track crossing behavior, using pairwise analysis of tracks from E6.5 to E7.0. Three unrelated pairs of nearby tracks during an E6.5 acquisition are shown in **G**. **H**. Track pair crossing events (each point is the mean of a half-embryo subset) are assessed in the anterior-posterior (top panel) and proximal-distal (bottom panel) axes, as a function of embryo stage. **I**. Quantification of division cohorts from E6.5 to E7.0 demonstrates separation behaviors of daughters following division (see **E**-**F** above).

Generally, *Mesp1* progeny filled the mesoderm layer in an orderly spatiotemporal pattern. Migrating from posterior regions, the nascent progenitors settled first in anterior and proximal locations, followed by progressively posterior and distal locations (Fig. 2B-C). We assigned 9 bins to the final destinations of the cells (after each 12-hour sequence), and analyzed the raw tracks for cell density, motility, and birthdate (Fig. 2B’). This showed that cells migrating within the posterior-distal locations were less dense, more motile, and born later than cells in anterior-proximal locations. In flat disc embryos such as those of most amniotes, this would be akin to an anterolateral-to-posteromedial sequence of mesoderm filling, guided by a concomitant density gradient.

SVF-processed tracks also demonstrated similar opposing gradients of birthdate and velocity (Fig. 2C). Quantitative analysis showed that extraembryonic mesoderm cells migrated more slowly than embryonic mesoderm cells (Fig. 2C’), consistent with prior findings [22]. However, as embryonic cells arrived in their positions and the mesoderm layer filled, they slowed to a velocity comparable to that of extraembryonic cells (Fig. 2C’).

Holistically, this process creates a dense pileup of slowing cells in anterior and proximal regions, juxtaposed with fast-moving sparser cells in distal and posterior regions that are still accumulating at their destinations. Thus, the embryo grows by posterior and distal extension (Fig. 2D’, S2B-C), similar to a traffic jam propagating along a highway, further and further from its origin. Embryos at 6.75 (late streak) exhibited similar opposing gradients of motility and density (Fig. S2D-F’), as nascent mesoderm is still being born at this stage. However, at E7.0, few new cells were born as embryos underwent ventral deformation and head folding (Fig. S2E).

Although the filling of mesoderm was orderly and stereotypical, we noticed that individual cell movements were quite chaotic during migration. This behavior has been observed qualitatively before [25,26,29], but our fluorescent reporters are uniquely suited for quantitative large-scale analysis of this phenomenon. Broadly speaking, gene expression and cell fate are patterned within the posterior epiblast and primitive streak [14,35], such that mesoderm and endoderm progenitors arise from distinct molecular and spatiotemporal regions. Yet, if an inflexible, precise proto-map within the mesoderm domain occurs prior to gastrulation, progenitors may be expected to migrate with mostly linear motion in order to preserve cell neighbor relationships and therefore the spatial map, as suggested by prior studies [14,22]. We observed the contrary.

Having manually tracked a large cohort of dividing cells during mesoderm assembly, we studied the migratory patterns of daughter cells (Fig. 2E-F), which necessarily share an ancestral site of origin in the primitive streak. To our surprise, daughter cells underwent substantial separation following division, up to 75μm (or 30% of total embryo length) in two hours (examples in Fig. 2E-F, 2I). When mother-daughter and daughter-daughter behavior were compared across stages, the strong separating movements of daughter cells declined as mesoderm assembly proceeded (Fig. 2I). By E7.0, daughters mostly remained in close proximity.

Similarly, we examined the crossing behavior of migrating cells during gastrulation using F-TGMM tracks, to determine the extent of mixing of unrelated cells. We observed frequent position swaps of co-migrating cells (Fig. 2G). By summarizing position swaps across two axes in various embryo regions, we again found that tracks cross each another less frequently with incrementing embryo stage (Fig. 2H). Comparing track pairs between E6.5 and E6.75 embryos, we likewise found tighter correlation of start and end positions in older embryos and in proximal (versus distal) locations (Fig. S2H-I’).

Collectively, these findings demonstrate that mesoderm assembly occurs in a stereotypical sequence from anterior-proximal to posterior-distal, guided by opposing gradients of density and cell motility. Moreover, considerable cell mixing occurs during this process, evidenced by the lack of preservation of cell neighbor relationships, until gastrulation finishes and positional settlement occurs.

### Birth of the *Smarcd3*-F6^+^ cardiac progenitors

Next, we examined embryos bearing the *Smarcd3*-F6-nGFP reporter, which utilizes a *cis* enhancer of BAF complex member *Smarcd3* / Baf60c termed “F6” that becomes active at E6.75 in cardiac precursors [6].

We empirically determined that nascent mesoderm at E6.75 could be grossly divided into two compartments on the basis of staining for either MSX1 or FOXC2 (Fig. 3A-B’), representing proximal and extraembryonic versus distal embryonic mesoderm, respectively. These populations likely correspond to distinct “destination cell types” in recent single cell RNAseq analysis of embryos at this stage [36], though they have not been spatially resolved heretofore. At E6.75, the earliest *Smarcd3*-F6^+^ progenitors definitively overlapped with the MSX1^+^ population (Fig. 3B-B’), but were distinct from cells expressing FOXC2 (Fig. 3A-A’). Since the *Smarcd3*-F6 lineage populates multiple tissues within all cardiac chambers [6], we next asked whether the *Smarcd3*-F6^+^/MSX1^+^ population is static or dynamic over time.

**Figure 3:**
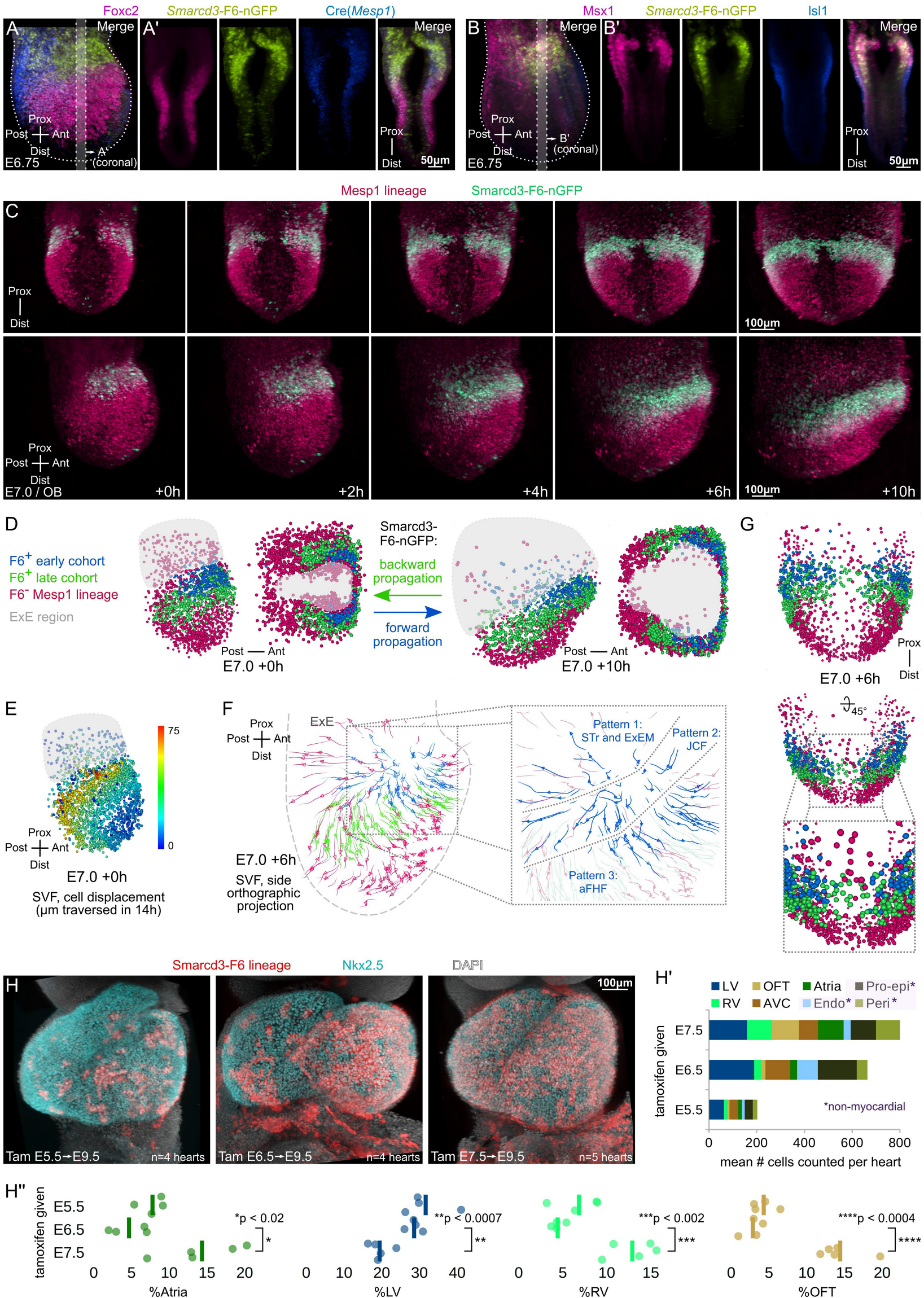
Birth of the Smarcd3-F6 cardiac progenitors. **A**-**B**. LSFM imaging of fixed E6.75 embryos shows that Mesp1-lineage-derived embryonic mesoderm is divided into two compartments, a distal FOXC2^+^;F6^-^ compartment (**A**), and a proximal MSX1^+^;F6^+^ compartment (**B**). **A’,B’**: Coronal slices from **A**,**B**. **C**. Time-lapse whole-embryo imaging at E7.0, demonstrating the onset and expansion of *Smarcd3*-F6-nGFP+ progenitors. Fused image sequence in frontal view (top row) and side view (bottom row). **D**. TGMM and SVF reconstruction of the total Mesp1-lineage population of cells, painted by F6 status: F6^-^, or either F6^+^ at the start (early cohort), or F6^+^ after twelve hours (late cohort). MaMuT display of the TGMM/SVF solution at the times indicated, from the side view (first and third panels in **D**), and Mercator projection (second and fourth panels in **D**). **E**. The same TGMM/SVF time-lapse sequence shown **in D** is depicted in MaMuT from the side view, with cells colored by the total displacement of their tracks. **F**. The TGMM/SVF solution is displayed in MaMuT (at the timepoint indicated) using an orthographic projection from the side, to depict the three classes of early F6^+^ cohort cells (inset). **G**. The midline breach forming the arch of the crescent is made up of both early- and late cohort F6^+^ progenitors. **H**-**H”**. Lineage tracing using *Smarcd3*-F6-CreERT2;Ai14 mice. NKX2.5 and DAPI counter-labeling as shown.

After E7.0, when the reporter was sufficiently bright for live imaging, an ongoing increase in *Smarcd3*-F6^+^ activity was apparent over the subsequent 12 hours (Fig. 3C, Video S2). However, dramatic ventral folding of the embryo become a moving target and obscured expression changes. Lateral mesoderm, especially, underwent greater apparent displacement than any other region during the sequence (Fig. 3E). We employed forward and backward propagation in SVF to mark F6^+^ tracks at the start (F6^+^ early cohort) and end (F6^+^ late cohort) of the sequence, respectively (Fig. 3D). Unexpectedly, we found that a large swath of mesoderm—much larger than the F6^+^ domain at E6.75—carried a cardiac fate. Thus, the F6^+^ domain expands distally (i.e. medially) and posteriorly as the reporter turns on, ultimately enveloping the *Smarcd3*-F6^-^/FOXC2^+^ domain seen at E6.75 (Fig. 3A). Indeed, the region with late reporter onset actually houses the majority of future cardiac progenitors (Fig. 3D).

A careful review of SVF tracks revealed that the early F6^+^ cohort had much greater migratory diversity than the late F6^+^ cohort, the latter of which moved outwardly and anteriorly following the overall expansion and ventral folding of the embryo (Fig. 3F). By contrast, we noted at least three patterns of early F6^+^ migration, including cells that: 1) migrated into extraembryonic structures; 2) traveled mostly posteriorly within the presumptive JCF space, laying on top of the forming crescent; and 3) followed the forming crescent (similar to the late F6^+^ cohort) anteriorly. Lastly, we noted that the anterior midline in the mouse was breached around E7.0 by lateral mesoderm bilaterally, and that these incursions across the midline were composed of both early and late F6^+^ cohorts (Fig. 3G).

When slightly later stage embryos (E7.0-E7.25) were examined, similar results were obtained (Fig. S3A-E), though the early F6^+^ cohort had already incorporated more distal and posterior regions by this point. Interestingly, expansion of F6 into more distal regions by E7.25 was paralleled by recession of FOXC2 and onset of ISL1 expression (Fig. S3F-F”). This suggests that distally (i.e. medially), the late F6^+^ cohort may be associated with SHF fate.

To concretely examine cell fate of the two cohorts, we used *Smarcd3*-F6-CreERT2 mice to lineage label progenitors at timepoints defined by tamoxifen administration. When tamoxifen was given at E5.5 or E6.5, we noted relatively similar contributions to myocardial structures, but with far fewer cells labeled at E5.5 (Fig. 3H-H”), consistent with the known onset of the reporter after E6.5. More interestingly, we noted a shift in the fates of E7.5-labeled cells toward SHF and outflow structures (Fig. 3H”).

While differential temporal fate of cardiac progenitors has been shown previously [7,14], it is significant here for two reasons. First, the graded onset of the F6 reporter (Fig. 3D) almost perfectly parallels the graded assembly of mesoderm by birthdate (Fig. 2C-D), except that it occurs 6-12 hours later. This parallel is further supported by the strikingly similar results of temporally-labeled *Mesp1* progeny [7] versus F6 progeny (Fig. 3H-H”): early *Mesp1*^+^ and F6^+^ cells contribute preferentially to LV, proepicardium, and AVC; late *Mesp1*^+^ and F6^+^ cells uniquely contribute to RV, OFT and atria. Second, the positions and marker co-expression of the two F6^+^ cohorts (as shown here) reveal patterning of the early cardiac crescent: anterolateral MSX1^+^ cells give rise to LV, proepicardium, and AVC, whereas posteromedial FOXC2^+^ cells, consistent with their apparent conversion to ISL1 expression (Fig. S3F-F”), contribute to RV and OFT. However, because the gradients of mesoderm accumulation and cardiac specification run diagonally in the embryo, spatially resolving late F6^+^ regions patterned for RV, OFT, or atria requires additional information [5,14], or tracking of later stage embryos.

### Mesenchymal-epithelial transition of the cardiac crescent

The next steps of cardiogenesis are not well studied in mammals, as tools for labeling the progenitors of interest and for examining their morphogenesis are scarce. Reporter mice such as those based on *Nkx2-5*, for example, initiate visible expression too late (E7.75 and beyond) to capture these stages [23]. Therefore, we again took advantage of *Smarcd3*-F6-nGFP reporter embryos from E7.25 to E7.75 to understand how the early crescent becomes suitable for forming a closed tube. During and after these stages, the pre-cardiac structures begin to take recognizable form [23], permitting annotation of patterned features and cell fates (Fig. 4A-B).

**Figure 4:**
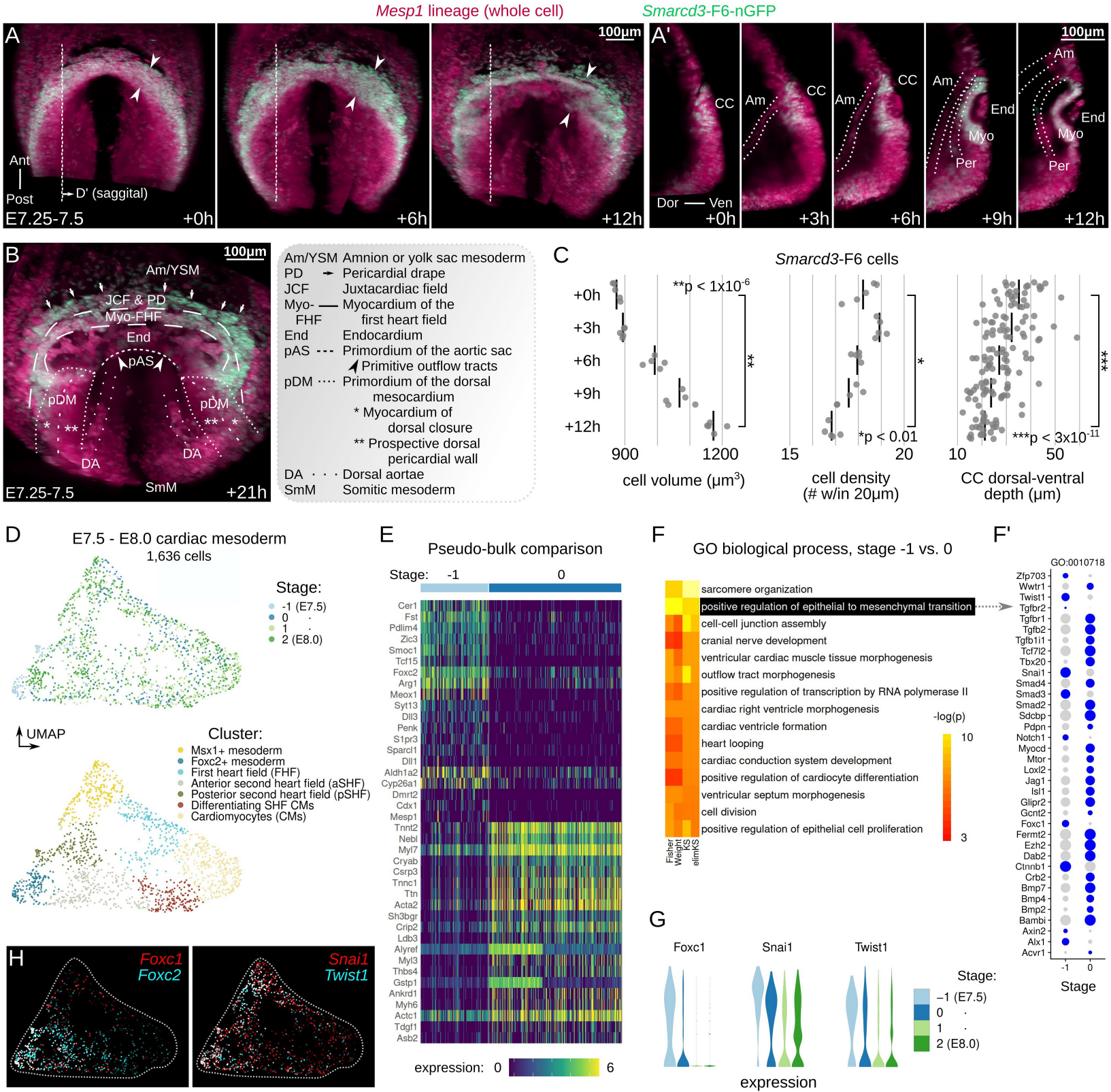
Mesenchymal-epithelial transition of the cardiac crescent. **A**-**B**. Time-lapse whole-embryo LSFM imaging at E7.25-E7.5 demonstrates splitting of the mesoderm and mesenchymal-epithelial transition of the cardiac crescent (arrowheads in **A**). Frontal (ventral) maximum projection (**A**) and single cutaway sagittal sections (**A’**). Am = amnion, Myo = myocardium, End = endocardium, Per = pericardium, CC = cardiac crescent. B. Annotation of early cardiac structures at +21h, **C**. Multi-modal estimations of *Smarcd3*-F6 cells during above sequence. **D**-**H**. Analysis of single cell RNA sequencing of early cardiac crescents at fine embryo stage resolution [9]. **D**. UMAP representation of earliest four stages of cardiac mesoderm. **E**. Differential gene expression between two earliest stages. **F**. Most significant 15 GO BP categories for significantly altered genes. **F’**. Differentially-expressed epithelial-to-mesenchymal transition genes. **G**. Violin plots of select EMT-governing transcription factors by timepoint. **H**. Overlap of select genes in UMAP space.

Despite the gross structural change of the cardiac crescent, the spatial expression of early cardiac-specifying transcription factors ISL1 and MEF2C remained static between E7.25 and E7.75 (Fig. S4A-C”). At first, the morphological changes (Fig. S4C’ and SC” vs. S4A’ and SA”, Video S3) appeared consistent with splitting of the mesoderm into splanchnic and somatic layers. However, in labeling by F6 (Fig. S4C-C” and 4A-A’) to reveal cardiac progenitor nuclei, we found a number of unexpected behaviors. First, the mesoderm simultaneously partitioned into three progeny compartments (Fig. 4A’): prospective endocardium, prospective myocardium, and prospective pericardium (i.e. somatic mesoderm). Second, the process appeared more complex than pure bisection (or trisection) of the mesodermal mesenchyme; the prospective myocardium flattened into a continuous single cell layer and expanded outwardly, stretching into the forming foregut pocket (Fig. S4C-C” and 4A-A’).

Next, we used a whole-cell tdTomato *Mesp1* lineage reporter to quantify the cells’ shape and size changes. Consistent with a transformation from dense mesenchyme to planar sheet, cell volume increased, cell density declined, and dorsal-ventral depth of the prospective myocardium decreased (Fig. 4C). Despite the subtle complexity, the morphological changes we observed (Figs. 4A, S4D, and S5A) are reminiscent of a mesenchymal-epithelial transition (MET), a critical morphodynamic step in numerous other developmental processes [37].

To investigate the possible mechanisms for a cardiac crescent MET, we analyzed single cell transcriptomes with fine temporal granularity during this process, dating from E7.5 to E8.0 [9]. Although mesoderm cells clustered principally by progenitor field and not by stage (Fig. 4D), we performed pseudo-bulk comparison between the two earliest stages (“-1” and “0”), i.e. E7.5 and intermediate between E7.5 and E7.75 (Fig. 4E). In scoring gene ontology (GO) biological processes (BP) for membership by differentially-expressed transcripts (Fig. 4F), we noted that the term “positive regulation of epithelial to mesenchymal (EMT) transition” was the second-most significant (Fig. 4F’). Almost universally, EMT regulatory transcription factors were downregulated at stage “0” compared with stage “-1,” and a panel of notable members—*Foxc1*, *Twist1*, and *Snai1*—showed clear temporal declines across the dataset (Fig. 4G). *Foxc1* was only present in the *Foxc2*^+^ population, whereas *Twist1* and *Snai1* were present in both mesodermal progenitor pools but declined during differentiation (Fig. 4H). These results identify a reversal of the transcriptional pathways taken for EMT, via down-regulation of positive EMT regulators, as a possible mechanism for cardiac crescent MET.

### Movement of cell populations during crescent MET

Because the observed MET occurs coincidentally with reshaping of the cardiac progenitor fields, together with spatial segregation of lineages (FHF, SHF, pericardium, endocardium, etc.), we next asked if we could reconstruct this process to determine patterns of cell fate and migration. With annotation guided by time-lapse LSFM footage from *Mef2c*AHF lineage tracing experiments (Figs. 5B, S5B-B”), we analyzed 9 tissues by backward propagation of the *Mesp1* lineage in SVF at E7.25 – E7.5 (Figs. S5A, 5A). Interestingly, pericardium and endocardium appeared to originate from progenitors interspersed within the cardiac crescent, yet they were spatially pre-configured within the mesenchyme by dorsal-ventral depth (Fig. 5A’), consistent with morphogen transfer between primitive germ layers [38].

**Figure 5:**
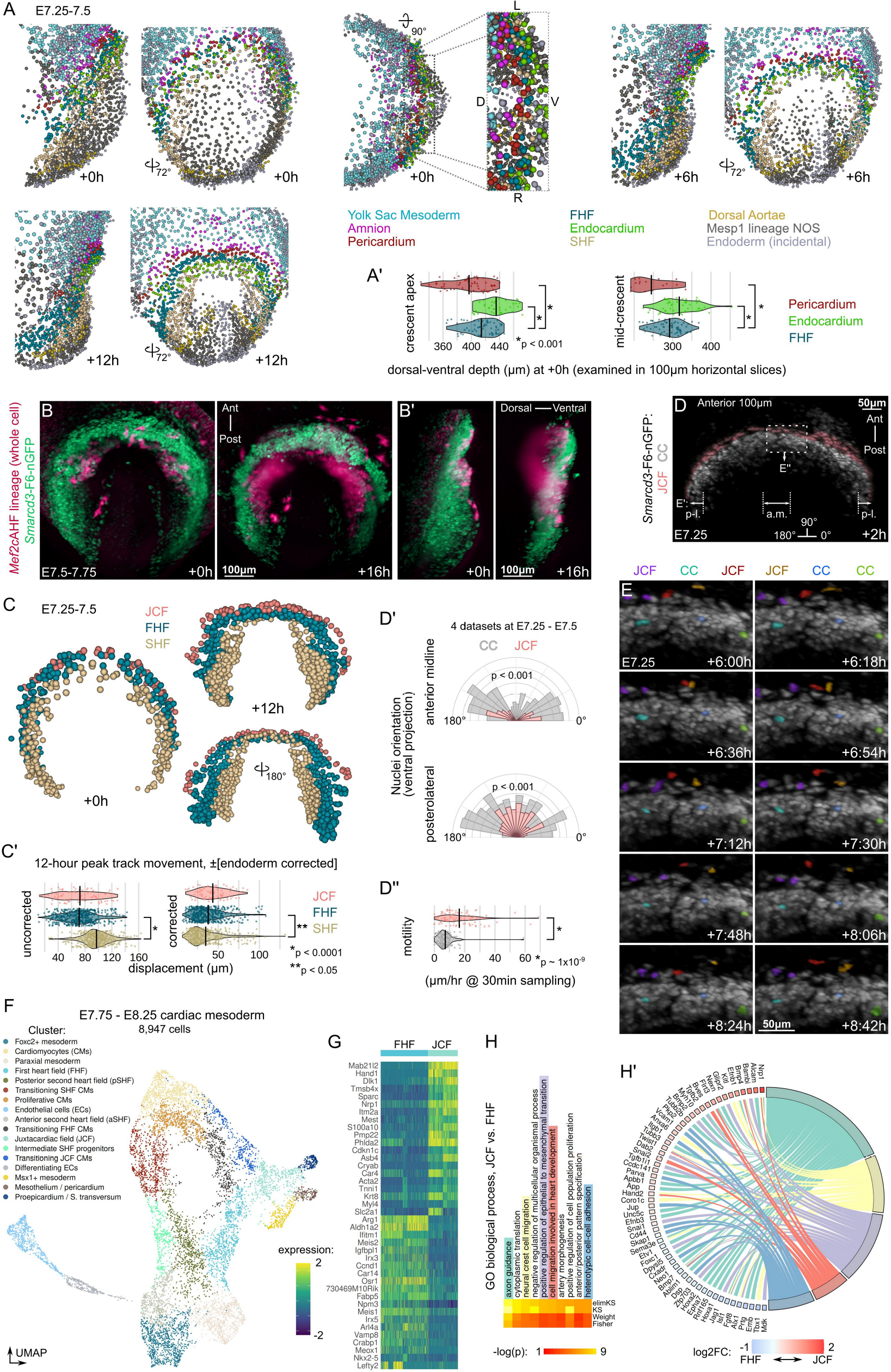
Movement of cell populations during crescent MET. **A**-**A’**. SVF reconstructions of cardiac crescent MET at indicated timepoints (**A**). Cell tracks, coordinates, and tissue assignments (color legend shown in **A**) are derived from backwards-propagated SVF data. **A’**. Positions of pericardial, myocardial, and endocardial cells in the dorsal-ventral axis determined through backwards propagation, for cells near the anterior crescent apex or in the middle of the crescent. **B-B’**. Time-lapse whole-embryo LSFM at E7.5-E7.75 using the *Mef2c*AHF lineage reporter, with ventral max projection (**B**) and lateral (**B’**) views. **C**-**C’**. SVF reconstruction at indicated timepoints from dorsal and ventral views, showing only FHF, SHF, and JCF. Peak SVF track displacement is quantified (**C’**, with endoderm correction in right panel). **D**. JCF cells are false colored in pink to highlight their position and orientation. **D’**. Quantification of long-axis orientations of nuclei comparing JCF to cardiac crescent (“CC”) cells (**D’**, top panel corresponds to anterior midline ‘a.m.’ in **D**, middle panel corresponds to posterolateral crescent ‘p-l.’ in **D**), with p-values from Watson U^2^ test. **D”**. Motility of JCF and CC cells. **E**. Time-lapse sequence from ventral partial max projection images using false colors to highlight a sample of six cells (3 JCF, 3 CC) throughout the sequence. **F**. UMAP space analysis of single cell RNA sequencing of mesoderm E7.75 and E8.25 [39]. **G**. Comparison of gene expression in FHF and JCF clusters; top log2FC differentially expressed genes plotted. **H**. Ten most significant GO BP categories for significantly altered genes. **H’**. Five interesting BP terms inspected by gene membership and log2FC differential expression (JCF versus FHF).

Next, we examined the myocardial fields, which expand considerably as they flatten into a one- or two-cell thick lamina. SVF propagation showed the myocardial fields formed a ventral ridge that extended dorsally into the deepening foregut pocket (Fig. 5C, Video S4). Although the SHF underwent greater movement, its net displacement was lower than either the JCF or FHF when corrected for endoderm deformation (Fig. 5C’), consistent with a coordination between myocardium and endoderm [16].

We next examined the JCF, which showed the greatest corrected SVF displacement of the three heart fields (Fig. 5C’). Curiously, we observed very brisk, seemingly chaotic movements (Fig. 5D and E, Video S4) within the JCF in all of our live experiments. JCF cells were F6^+^, ISL1^-^, and had varying levels of MEF2C (Fig. S5D), and JCF nuclei were more tangentially oriented along the crescent than FHF and SHF (cardiac crescent, “CC”) nuclei (Figs. 5D’ and S5C). Consistent with visual observations, JCF cells were far more motile than the relatively immobile CC cells (Fig. 5D’ lower panel and 5E).

To investigate gene expression that could be responsible for this behavior, we compared the JCF and FHF (Figs. 5F-H’, S5E-G) by single cell RNAseq [39]. In scoring GO BPs for membership by differentially-expressed genes between JCF and FHF (Fig. 5H), we found a number of significant terms that incorporate motility, adhesion, or migration (Fig. 5H), and plotted a collection of differentially expressed member genes (Fig. 5H’). *Nrp1*, a member of three such GO BP terms, was the most upregulated in the JCF versus FHF. *In addition*, a number of bone morphogenic protein and other matrix/guidance molecules were differentially expressed in the JCF (Fig. 5H’).

### Transformation of the epithelial cardiac crescent into the early heart tube

Shortly after becoming an epithelium (E7.75), cardiac progenitors undergo rapid morphogenesis to form (E8.0) and dorsally close (E8.25) the early heart tube [23]. Using in toto LSFM imaging with TGMM/SVF reconstruction at E7.75 (Fig. 6A-A”, Video S5), we observed two notable patterns of movement. First, progenitors within the dorsal aspect of the epithelial sheet lifted off the endocardial surface, causing the ventral-anterior portion of the ridge (along with the overlying JCF) to rotate posteriorly (pattern 1, arrowhead in Fig. 6A’) by pivoting on FHF/JCF boundary, which acted as a morphological anchor. The JCF followed the ventral torsion of the ridge, being dragged and nearly draped around the ventral aspect of the cardiac epithelium. Second, a knob-like epithelial protrusion propagated posteriorly within the dorsal aspect of the crescent, traveling posteriorly as a wave (Fig. 6A and pattern 2, arrowheads in 6A”).

**Figure 6:**
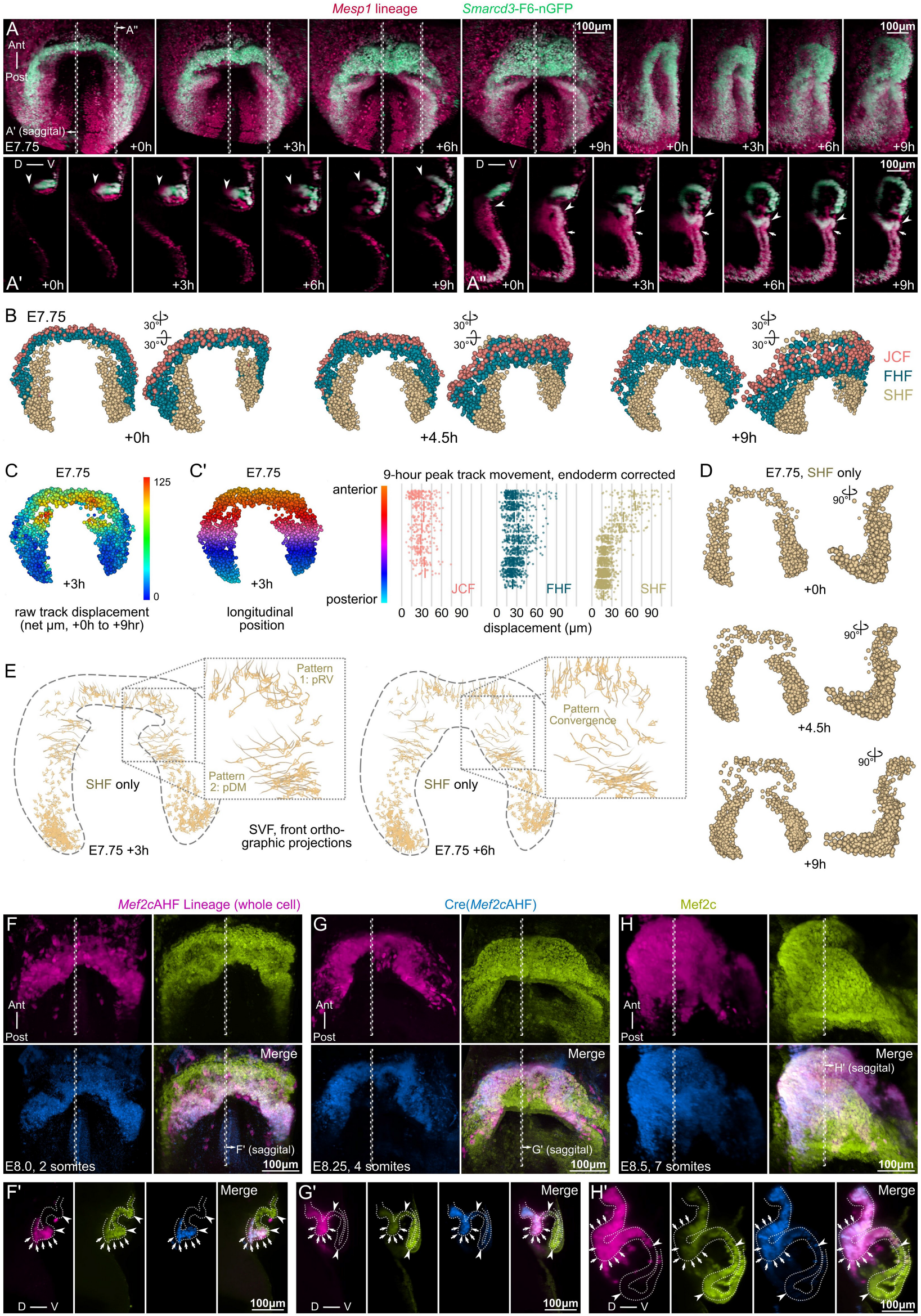
Transformation of the epithelial cardiac crescent into the early heart tube. **A**-**A”**. Time-lapse whole-embryo LSFM imaging starting at E7.75, with max projection ventral (left four panels in **A**) and lateral oblique (right four panels in **A**) views. 7.5μm thick sagittal slices from indicated regions in **A** are also shown (**A’**, **A”**). Arrowheads and arrows in **A’** and **A”** point to congruent cells at different timepoints. **B**. SVF reconstructions of tracked images series are shown at indicated timepoints from ventral and angulated views, with only FHF, SHF, and JCF cells drawn. **C**-**C’**. Quantitative analysis of SVF reconstruction shown in **B**, with cells colored by their track’s displacement (**C**) or by their anterior-posterior position (left panel in **C’**). **D**. SVF reconstructions were re-drawn from ventral and lateral views, with only SHF cells at indicated timepoints. **E**. Examination of morphogenic dynamics within the SVF revealing two distinct regional patterns. **F**-**H’**. LSFM examination of fixed embryos for lineage tracing of *Mef2c*AHF, during LHT formation. Max projection ventral views labeled for MEF2C protein, *Mef2c*AHF-Cre, and *Mef2c*AHF lineage are shown at 2 somites (**F**), 4 somites (**G**), and 7 somites (**H**). Midline sagittal planes (7.5μm thick slices) at indicated stages (**F’**, **G’**, **H’**) are examined for movement of SHF cells into the heart, a process that leads to dorsal mesocardium formation and closure of the LHT.

SVF reconstructions of this sequence, annotated using time lapse imaging of *Mef2c*AHF lineage tracing experiments (Fig. 5B and S5B-B”), indicated that these two patterns were features of the SHF (Fig. 6B-C). Quantitative tracking showed that SHF cells underwent much greater displacements than either JCF or FHF (Fig. 6C’) during this process. Lastly, review of orthogonal SVF projections revealed the anterior-(pattern 1) and medial-(pattern 2) directed torsional motion resulted in opening and closing, respectively, of the early heart tube (Fig. 6D and left panel in 6E). As the midline was reached by prospective dorsal mesocardium / dorsal closure myocardium, those patterns converged to drive the epithelial sheet anteriorly into the forming heart (Fig. 6E, right panel).

In fixed embryos labeled with MEF2C^+^ (JCF, FHF, and SHF) and *Mef2c*AHF lineage (SHF only) cells, we found that SHF progenitors entered the forming linear heart tube (LHT) via wave-like translocation or treadmilling of the SHF epithelium through the knob-like structure (Fig. 6F’, G’, H’, arrowheads demarcate FHF boundary, arrows point to dorsal wave). Taken together, these experiments shed light on the diverse morphodynamics of the SHF, both in space and time [40], indicating that they concurrently enact dorsal closure, formation of dorsal mesocardium, and establishment of the arterial pole (see next section).

### LHT closure by *Isl1*-dependent morphogenetic wave within differentiating SHF progenitors

Empirical attempts to characterize the SHF epithelial knob-like structure revealed that it was labeled by intermediate expression of ISL1 and NKX2-5 (Fig. S6A-A”’). By single cell RNAseq analysis, this zone (intermediate *Isl1* and *Nkx2-5*) resolves to the “Transitioning SHF CMs” cluster, for which a key marker gene was *Tdgf1* (Fig. S6B-B’). The unique molecular features of the knob also include a number of extracellular signaling and cytoskeletal factors (Fig S6C-F). Future investigation into these may shed light on the dramatic morphogenetic behaviors of the knob, and formation of SHF structures.

Next, we examined *Nkx2-5* or *Isl1* mutant embryos at E8.5, when dorsal seam myocardium had reached the midline in control LHTs (Fig. S6G”, see arrowheads). The comparable region in *Nkx2-5* mutant embryos appeared disorganized, over-folded, and delayed in its approach to the midline (Fig. S6H”, see arrowheads). *Isl1* mutants, on the other hand, failed to form the knob/wave region entirely (Fig. S6I”, see arrowheads), and therefore retained an open configuration of the prospective dorsal mesocardium and aortic sac. We speculate that this is due to a paucity of SHF cells from decreased proliferation in the SHF [12,41], or more likely from mis-specification of early *Foxc2*^+^ progenitors to non-cardiac fates [42]. Indeed, LSFM analysis of later stage *Isl1* KO embryos revealed unusual co-expression of FOXC2 and TNNT2 (Fig. S7A’ versus S7B’), as well as complete absence of dorsal LHT closure (Fig. S7A” versus S7B”).

### Loss of *Mesp1* disrupts the density gradient that forms after gastrulation, altering mesoderm organization

To gain an understanding of cues that control the spatiotemporally-governed early cardiac progenitor behaviors, we studied gastrulation in *Mesp1* mutants, where early organogenesis does not occur due to specification and/or migration defects [6,8,36,43]. Although the movement behaviors of *Mesp1* knockdown cells have been studied *in vitro* [44], we exploited LSFM and our tracking workflow to better understand their actions *in vivo*.

We observed that the anterior flank of *Mesp1* KO mesoderm did not reach the anterior midline (Figs. 7A-A” versus 7B-B”, Video S6). F-TGMM tracks from these LSFM experiments were grouped/binned by their destination position along the anterior-proximal to posterior-distal filling gradient we previously determined (Fig. 7C-C”). Track birthdate and motility gradients were preserved in mutants (Fig. 7C’, top and middle panel pairs), but the density gradient appeared flattened and partially inverted (Fig. 7C’, bottom panel row). When we examined the spatial vectors of these tracks, binning the gradient into three sections, we observed a severe defect in anterior-directed, as well as outward expansive motion (Fig. 7C”). To explore possible mechanisms for failed directional migration, we analyzed *Mesp1* KO embryos by single cell RNAseq (Krup et al., manuscript in preparation). A host of morphogens, their receptors, and downstream signaling effectors [45] are perturbed in *Mesp1*-null mesoderm progenitors. This includes *Rac1* and *Fgf* genes, which have been shown to be important for directed motility of the mesoderm [22,45].

**Figure 7:**
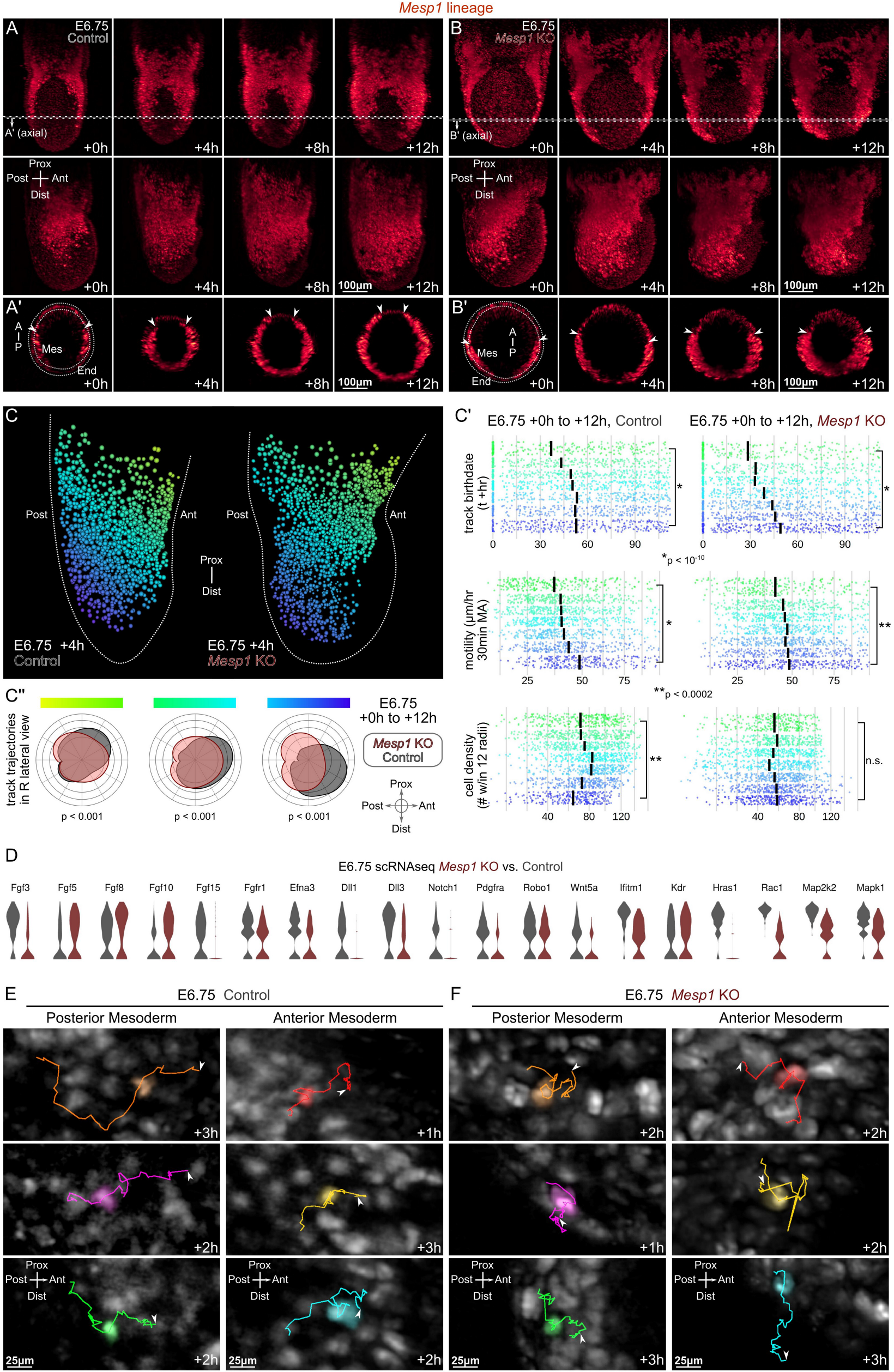
Loss of *Mesp1* disrupts the density gradient that forms following gastrulation, altering mesoderm organization. **A**-**B’**. Time-lapse whole-embryo LSFM imaging at E6.75. Control embryos: **A**-**A’**; *Mesp1* mutant embryos: **B**-**B’**. Max projection views are shown from ventral view (top row in **A** and **B**) and lateral view (second row in **A** and **B**). 7.5μm-thick axial cutaways from indicated regions in **A** and **B** are also shown (**A’**, **B’**). Mes = mesoderm, End = endoderm, A/Ant = anterior, P/Post = posterior, Prox = proximal, Dist = distal. **C-C’**. Quantitative analysis of raw TGMM tracks in control (left panels in **C-C’**) and *Mesp1* mutant (right panels in **C-C’**) time lapse data. C”. Density distribution of mesoderm track trajectories. **D**. Single cell RNA sequencing of gastrulating mesoderm progenitors, with various migration-related features depicted by expression. **E**-**F**. Qualitative analysis of tracks (of duration 4-6h shown) in control (**E**) and *Mesp1* mutant (**F**) time lapse series. Left panel column in each of **E** and **F** show TGMM tracks originating in posterior regions, where right panels in each show anterior originating tracks. Arrowheads in **E** and **F** demarcate track endpoints.

Examination of individual tracks in *Mesp1* KO embryos revealed near absence of anterior-directed motion in the anterior, older-born cells (Fig. 7F versus 7E), whereas younger, posterior cells maintained some degree of anterior movement. This abnormal movement of older-born cells may underlie the observed density gradient inversion, as it leads to accumulation of cells in the middle of the embryo rather than antero-proximally (Fig. 7B’ vs. 7A’). Directionality, not motility, may thus be the culprit for disorganized *Mesp1* mutant mesoderm.

## Discussion

In this study, we first aimed to overcome the big data intimidation of LSFM, allowing us to focus on several deep biological questions concerning early cardiac fate and morphogenesis. Our improved comprehensive workflow is an important step to simplify and democratize the complexities of live LSFM. Its software components are open source, portable, and easier to use than ever before, and the requisite hardware is accessible to non-microscopists (such as ourselves). Armed with fluorescent reporter mice, a widely available LSFM instrument, and this computational toolbox, we embarked on a study spanning a short yet dramatically important window in mouse development, from gestational days 6.5 to 8.0 (Fig. S7D). We identified distinct patterns of mesoderm filling, multipotent cardiac identity, and morphogenesis that critically underlie the emergence of the LHT.

During gastrulation, we observed opposing gradients of progenitor density and motility, similar to the manner in which a traffic jam propagates along the highway further and further from its origin. In presomitic mesoderm of chicks, a random motility gradient controls axis extension [46,47], and our observations of mouse lateral plate mesoderm are qualitatively and quantitatively similar. Somites, however, are periodic structures, whereas the heart is a singular object formed from the collective migration of precursors to a final destination. Therefore, the motility of lateral mesoderm is unlikely to be completely random, even if it appears quite chaotic. Our examination of *Mesp1* mutants clearly portrays initial mesoderm migration as directed, as *Mesp1* KO embryos do not form a density gradient, and cells lose directionality, thus preventing the completion mesoderm assembly.

Gastrulating zebrafish embryos are resilient to cell mixing, utilizing morphogenetic gradients to ultimately establish mesoderm patterning [48]. Our analysis demonstrates re-arrangement and crossing of cell tracks during gastrulation in the mouse, suggesting that considerable early plasticity must exist among progenitors with respect to final cell fate. Thus, it seems unlikely that rigid fate allocations are present in the primitive streak region before gastrulation, but instead that general trends (i.e. MSX1^+^ vs. FOXC2^+^ nascent mesoderm) are followed with flexibility in each embryo. This reinforces the necessity of patterning cues or landmarks, largely established through morphogen gradients, that are present in the early embryo during and after arrival of mesoderm [49,50]. The distinct spatial patterns of mesoderm assembly and heart field specification, which are oblique to one another (Fig. S7C), likely explain the observed pre-configuration of cardiac fates at the time of gastrulation.

As gastrulation terminates, we demonstrate that myocardial progenitors undergo a previously unappreciated mesenchymal-epithelial transition (MET) that rapidly extends throughout the entire cardiac crescent. Moreover, we identify a temporal reduction of EMT gene expression that may have a role in this process. Further interrogating these regulators, such as through co-expression network analysis (Fig. S4H), may spark future endeavors to understand mesodermal and cardiac MET. From a structure/function perspective, efficient cell filling of the originally-empty mesoderm layer benefits from free-form movement of progenitors, guided by each other and surrounding cues [45,51]. However, formation of the heart requires each cell’s movement to act upon all others in the crescent, so that net morphogenesis becomes an emergent property of the collective – and here, an epithelium is well-suited.

Once the cardiac epithelium is forged, regional discrepancies in morphogenetic behaviors emerge, such as the brisk dance-like movements of the JCF. Our data do not provide a teleological reason for the motility of the JCF. However, it may be that JCF cells are not constrained until the torsional movement of the FHF ridge and push from the SHF drives them into their pro-epicardial alcove. In terms of the SHF, whose anterior ‘pushing’ behavior is necessary for opening of the early heart tube, a dramatic wave of differentiation and morphogenesis actually propagates posteriorly. This pull and push mechanism is unanticipated, and provides a new insight into the formation of the LHT. Moreover, dramatic medial extension of the SHF forms the heart’s dorsal closure and concurrently separates inflow and outflow myocardium.

Combining these data, we establish a holistic model of early cardiogenesis to unify these findings and reconcile prior evidence (Fig. S7C). Here, the prospective LV (and pro-epicardium) lies at the farthest anterior-lateral extent of the crescent, which is the earliest born during gastrulation. Immediately medial to the LV lies the prospective RV. Within the RV progenitors, the *Tdgf1*^+^ knob forms as RV and ultimately OFT progenitors are incorporated into the heart. Lateral to the dorsal closure lie the atrial progenitors, which are pulled anteriorly within the epithelial sheet, adding to the venous pole. Still, many unanswered questions remain, such as the necessity and sufficiency of these morphodynamics to heart formation, and the complex molecular events that trigger such dramatic activity.

Overall, our results shed light on an obscure window in early mammalian development, by connecting discrete morphological events in sequence with fine spatial and temporal resolution. This illuminates individual and collective movements of mammalian organ precursors, their origin and dynamic spatial relationships, and the complex and carefully choreographed morphogenetic steps in the formation of an embryonic organ. These findings make an important contribution to cardiac-specific, but likely generalizable features of cell allocation, which may ultimately be identified as broad themes in embryogenesis.

## Acknowledgements

We thank Blaise Ndjamen for help with microscopy, and Kathryn Claiborn for editorial guidance. This work was funded by a grant from the NHLBI (R01 HL114948) and The Younger Family Fund. M.H.D. was supported by NIH T32 training grants 2T32-HL007731-26 and T32-HL007843-24, as well as funding from UCSF Department of Medicine, Division of Cardiology. A.L.K. was supported by scholarships from the NSF (GRFP 2034836) and the AHA/CHF (817268). J.M.M. was supported by an NIH T32 training grant 5T32-HL007544-34, and NIH F32 fellowship 1F32-HL162450-01. This work was also supported by an NIH/NCRR grant (C06 RR018928) to the J. David Gladstone Institutes.

## Author Contributions

M.H.D. and B.G.B. designed the project. M.H.D. imaged all live embryos, performed immunostaining and imaging, developed computational tools, and analyzed the data. A.L.K. generated scRNAseq data. J.M.M. benchmarked quantitative analyses. M.H.D. wrote the paper with input from the other authors.

---

**Figure S1:**
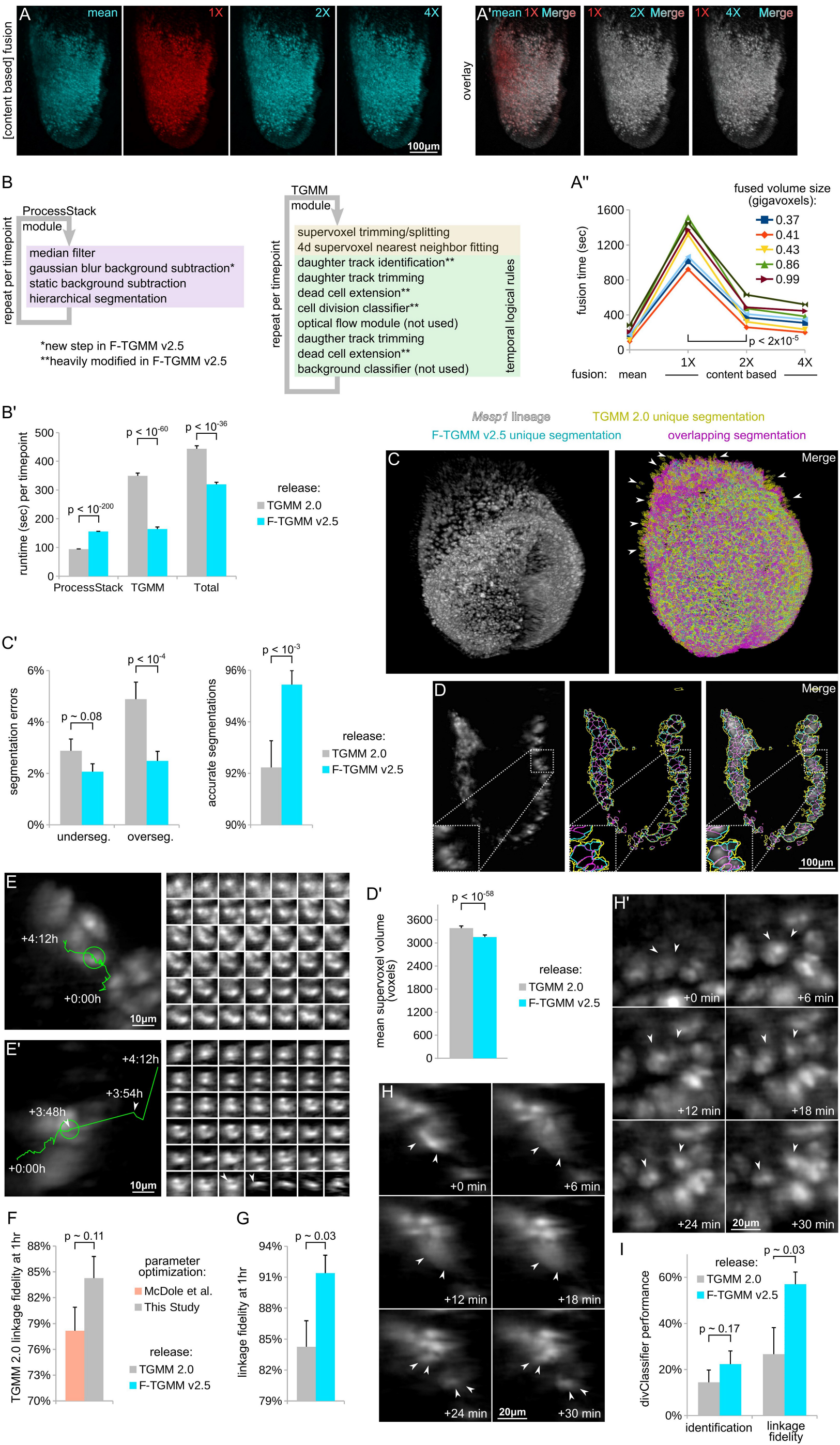
Improvements to LSFM computational workflow, comparative analysis, related to Figure 1. **A-A”**. Speed-optimized (2X or 4X downsampled weight) content-based fusion produces nearly identical results as the original content-based fusion algorithm (1X), but with much lower computational overhead. Comparisons of embryo max lateral projections (**A**) show qualitative benefit to content-based rather than mean fusion (left panel in **A’**). 2X and 4X speed optimized methods yield similar benefit (second and third panels in **A’**). Comparisons of CPU time for the various methods across several datasets is shown in **A”**. **B**-**B’**. Refinements to the main loops of TGMM components, yielding improved overall tracking efficiency for F-TGMM v2.5 compared with TGMM 2.0. Benchmarks of component runtimes is shown in **B’**. **C-D’**. Segmentation accuracy of F-TGMM v2.5, showing oversegmentations (arrowheads in **C**) and undersegmentations (inset in **D**) that are corrected by dynamic background subtraction. Segmentation errors and total accurate segmentations are quantified in **C’**. Also due to dynamic background subtraction, mean supervoxel size is lower in F-TGMM v2.5 (**D’**), which contributes to the overall improved efficiency of F-TGMM v2.5 versus TGMM 2.0. **E**-**G**. Assessment of tracking. Two 4-hour cell tracks are shown, one with correct linkage across the entire timespan (**E**), the other with a two large shifts (**E’**, at +3:48h and +4:06h) that reflect incorrect linkage to a distant, unrelated cell. In **F**, tracking accuracy in 1-hour segments was quantified for TGMM 2.0, using empirically optimized parameters versus those previously published as optimal [29]. In G, tracking accuracy in 1-hour segments was quantified for TGMM 2.0 vs. F-TGMM v2.5. **H**-**I**, Division accuracy quantification of TGMM 2.0 versus F-TGMM v2.5. In **H**, a correctly tracked division event with mother and daughter cells (arrowheads in **H**) shown in time lapse. In **H’**, an incorrect cell division occurred when two neighboring cells (arrowheads in **H’**) separated. **I**. Quantification of cell division tracking in TGMM 2.0 versus F-TGMM v2.5, by primary identification of cell divisions and by linkage accuracy once a division is identified.

**Figure S2:**
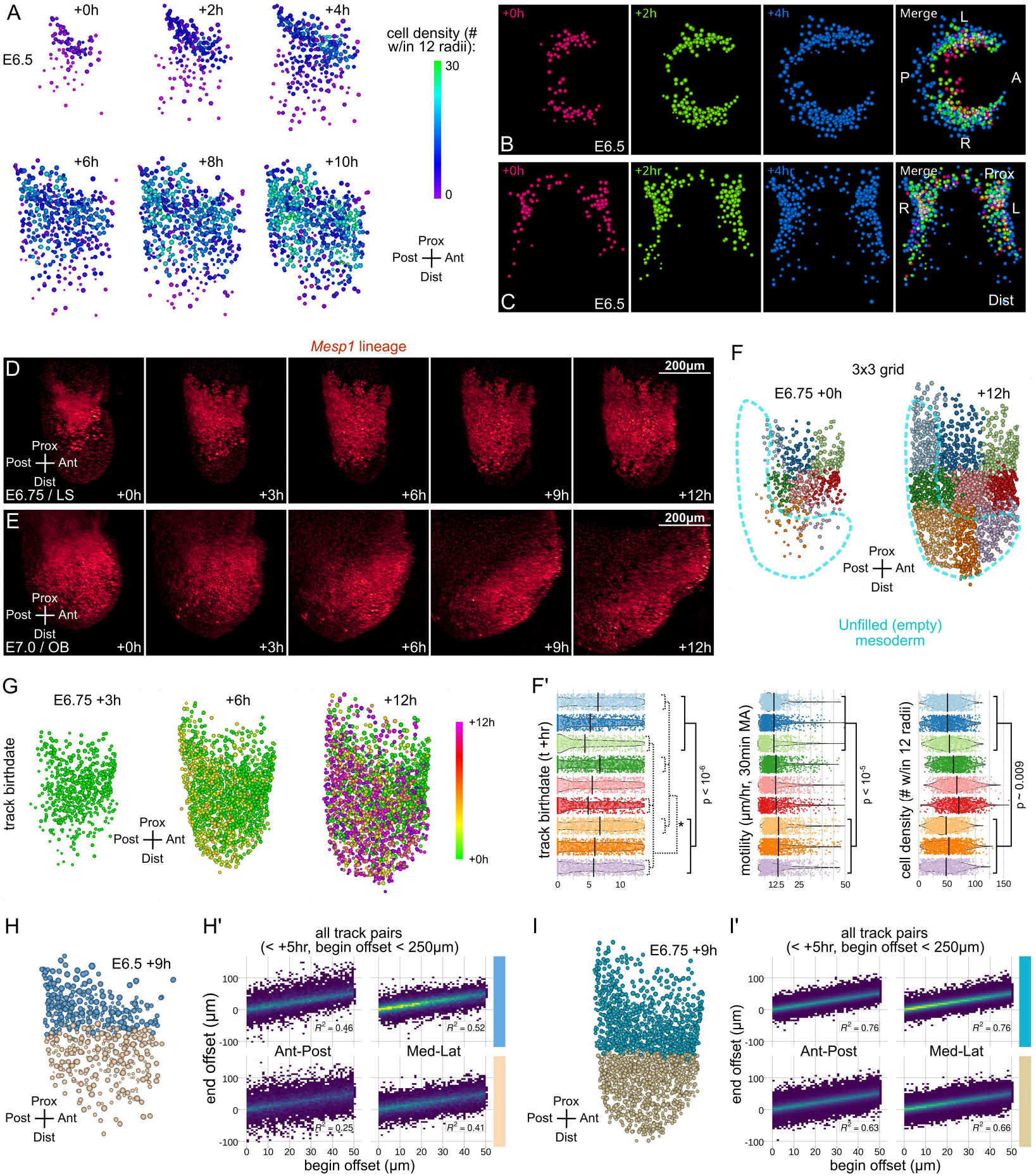
A spatiotemporal gradient of mesoderm accumulation, related to Figure 2. **A**. TGMM reconstruction of E6.5 embryo live imaging, cells painted by cell density. **B**-**C**. TGMM/MaMuT reconstructions of E6.5 mesoderm filling, as seen from top (**B**) and front/ventral views (**C**). Cells are uniformly colored by timepoint snapshotted (with time-lapse merge in the right-hand panels). **D**-**E**. Time-lapse whole-embryo LSFM imaging of *Mesp1* lineage at E6.75 (**D**) and E7.0 (**E**), showing right-half max Z-projections. **F**. Side views of a TGMM/MaMuT reconstructed E6.75 embryo during live imaging, with all tracks retrospectively partitioned onto a 3×3 grid (anterior-proximal in light green, posterior-distal in lilac). Spatial binning shows mesoderm filling of newborn cells into progressively posterior and distal/medial regions. **F’**. TGMM tracks, in each 3×3 bin (also including the nearest half of tracks belonging to each adjacent bin) were analyzed from +0h to +15h for birthdate (track start timepoint), motility (average of moving average velocity sampled over 30-minute window), and cell density (number of other cells within 12 radii of each). **G**. Side view TGMM reconstruction of E6.75 embryo during live imaging, with cells painted by track birthdate. **H**-**I’**. In **H** (E6.5) and **I** (E6.75), cells are divided into two bins based on their proximal-distal position. All pairs of TGMM tracks originating during acquisitions from +0hr to +5hr are considered, if the cells are initially offset by less than 250μm in the specified axis (proximal-distal or anterior-lateral) of analysis. End offset is plotted as a function of begin offset (E6.5 in **H’** and E6.75 in **I’**) in a density scatterplot, with linear correlation coefficient shown. Note the greater correlation (i.e. greater correspondence in the position of each track pair) in proximal positions and E6.75 versus E6.5.

**Figure S3:**
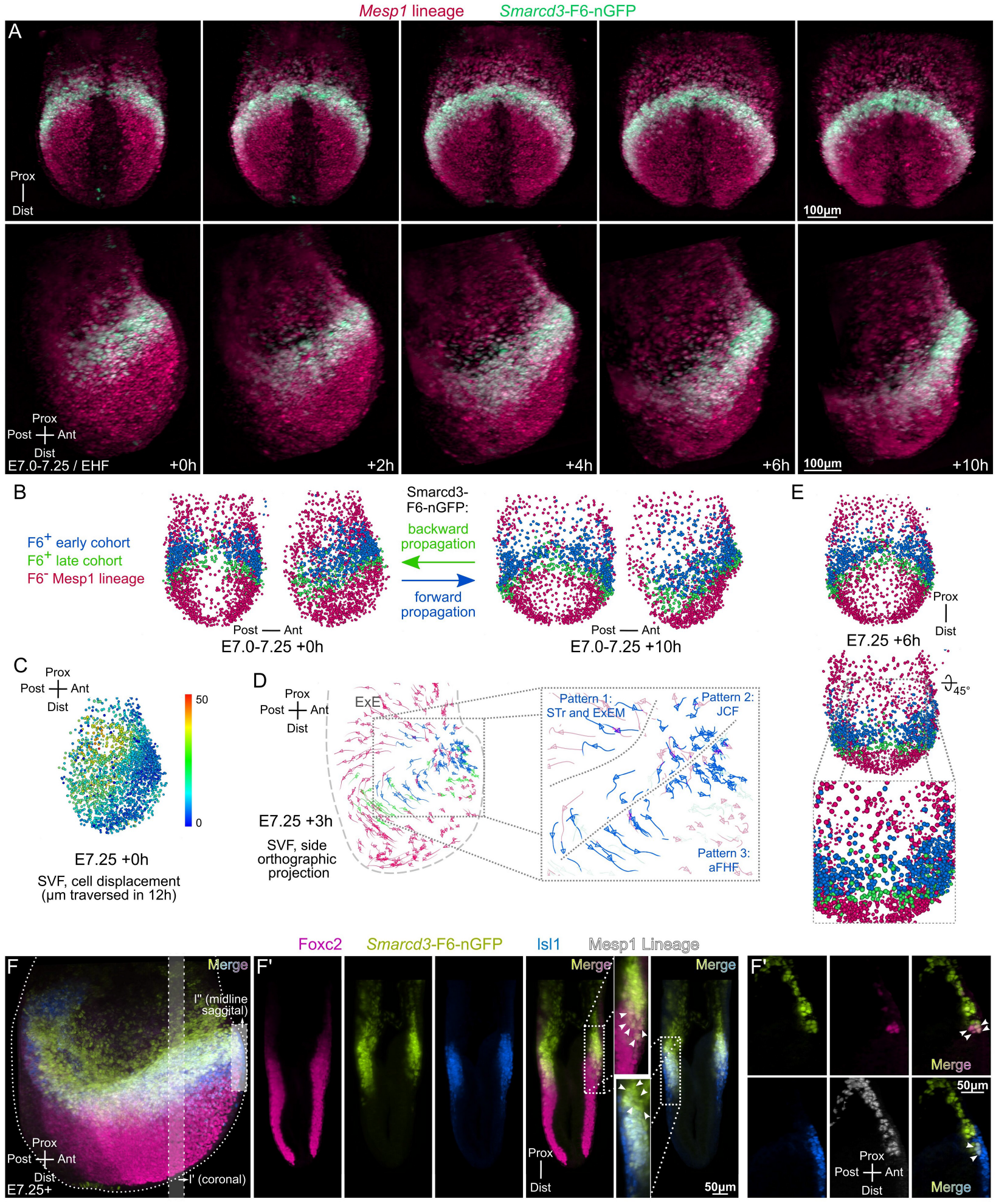
Birth of the Smarcd3-F6 cardiac progenitors, related to Figure 3. **A**. Time-lapse whole-embryo imaging by light sheet microscopy at E7.0-E7.25, an alternate dataset that demonstrates the onset of cardiac specification (*Smarcd3*-F6-nGFP). The fused image sequence is displayed from the frontal (ventral) view (top row) and side view (bottom row). **B**. TGMM and SVF reconstruction of the dataset shown in **A**, with the total *Mesp1*-lineage population painted by F6 status (colors as indicated): F6^-^, or either F6^+^ at the start (early cohort), or F6^+^ after ten hours (late cohort). MaMuT display of the TGMM/SVF solution at the times indicated, from the front/ventral view (first and third panels in **B**), or side view (second and fourth panels in **B**). **C**. The same TGMM/SVF time-lapse sequence shown above is depicted in MaMuT from the side view, with cells colored by the total displacement (units and scales as shown) of their tracks. **D**. The TGMM/SVF solution is displayed in MaMuT (at the timepoint indicated) using an orthographic projection from the side, to depict the three classes of early F6^+^ cohort cells (inset) by migration pattern. **E**. The arch of the crescent is made up of both early- and late cohort F6^+^ progenitors, as seen from the front (ventral) view in MaMuT (top panel in E), and further confirmed in zoom/rotation (middle panel in **G** and inset). **F-F”**. By E7.25, *Smarcd3*-F6 labeling has spread distally and posteriorly (side view max projection in I), and coronal sections show it now encompasses FOXC2^+^ early mesoderm progenitors (first merge panel from the left in **F’**). Also at E7.25, ISL1 is present in F6^+^ cardiac progenitors, but is excluded from prospective JCF cells overlying the forming crescent (second merge panel from the left in **F”**). At the now-crossed ventral anterior midline (narrow partial side view projection in **F”**), F6^+^;ISL1^-^ and F6^+^;FOXC2^+^;ISL1^+^ progenitors are present, corroborating findings from TGMM/SVF and lineage tracing experiments above.

**Figure S4:**
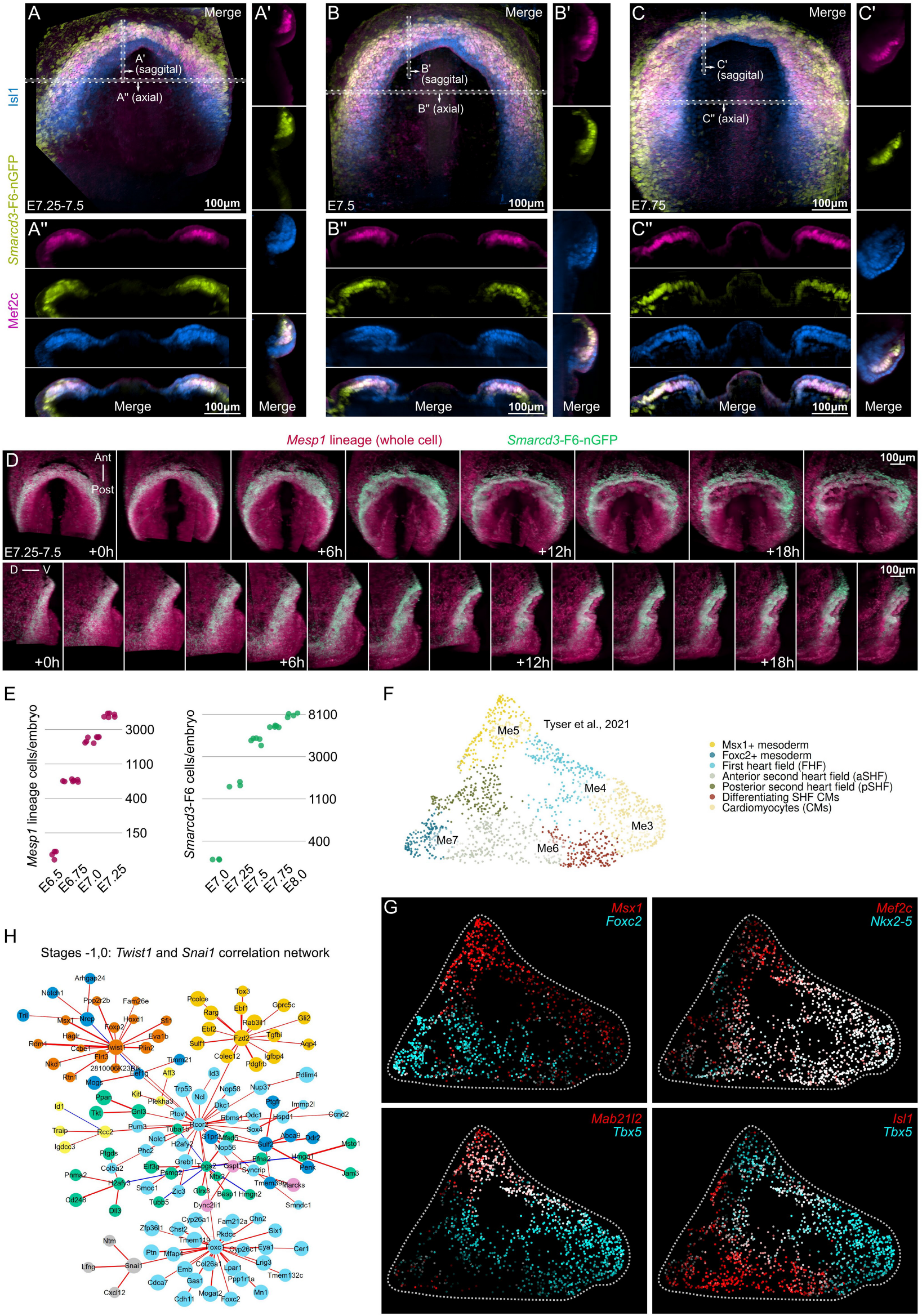
Mesenchymal-epithelial transition of the cardiac crescent, related to Figure 4. **A-C”**. LSFM imaging of fixed E7.5-E7.75 embryos shows that the cardiac crescent transforms dynamically during foregut pocket involution. Ventral maximum projections (**A**,**B**,**C**) demonstrate subtle morphological changes within the cardiac crescent. Sagittal slices in the arch of the crescent are shown (**A’**,**B’**,**C’**), as well as axial slices (**A”**,**B”**,**C”**), showing a similar transformation to flat sheet. **D**. Time-lapse whole-embryo imaging by light sheet microscopy at E7.25-E7.5, demonstrating splitting of the mesoderm and mesenchymal-epithelial transition of the cardiac crescent. The fused image sequence is displayed from the frontal (ventral) and lateral partial maximum projections. **E**. Estimated *Mesp1* lineage (based on TGMM cells in each frame of *Mesp1* lineage tracked datasets, accounting for an estimate of endoderm and non-bona fide mesoderm) and *Smarcd3*-F6 (based on TrackMate detection, with DoG radius 15px and threshold between 6 and 7) cell counts at stated timepoints across several embryos. **F**. UMAP shown in Fig. 4D, with superimposed labels corresponding to the clusters as named in source publication [9]. **G**. Notable markers are depicted in qualitative co-expression feature plots. **H**. Pearson correlation is performed on Twist1 and Snai1 across all features and cells of the dataset, and the correlation matrix for the top 70 candidates with strongest coefficient for each is used to construct a speculative gene regulatory network, using a |r| > 0.25 cutoff. Blue lines indicate negative correlations, edge width indicative of |r|.

**Figure S5:**
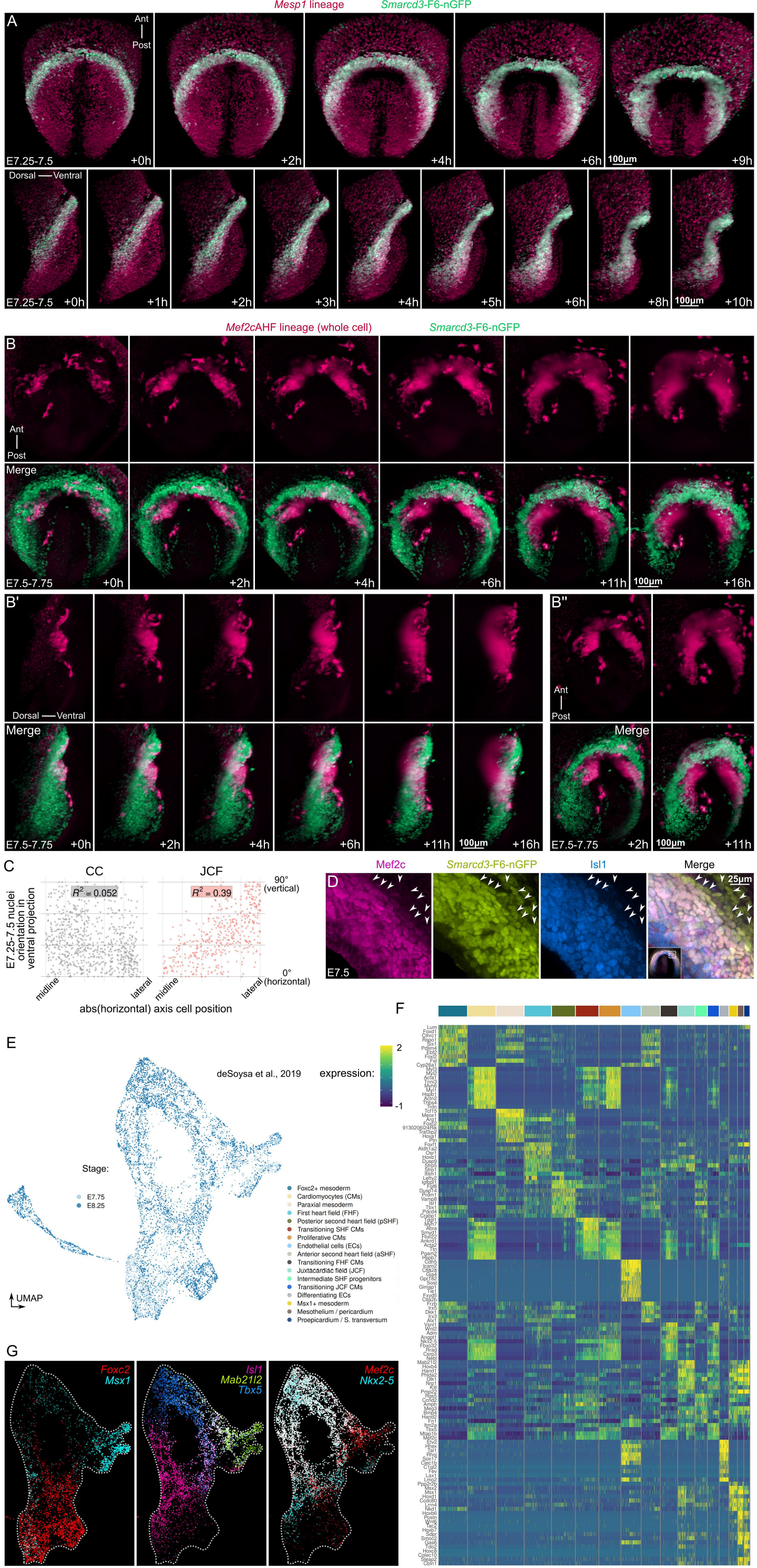
Movement of cell populations during crescent MET, related to Figure 5. **A**. Time-lapse whole-embryo light sheet imaging starting at E7.25 – E7.5, revealing mesoderm splitting and the cardiac crescent MET. Top row is ventral maximum projection, bottom row is lateral partial maximum projection. The image sequence is segmented, tracked, and SVF reconstructions are analyzed in the accompanying main figure. **B-B”**. Time-lapse whole-embryo light sheet imaging starting at E7.5 – E7.75, demonstrating the onset of second heart field identity, morphogenesis of the prospective aortic sac region, and formation of the prospective dorsal mesocardium. Ventral maximum projection views are shown (**B**), as well as lateral partial maximum projection (**B’**) and oblique frontal maximum projection (**B”**). **C**. Quantifications of JCF versus CC long-axis orientations of nuclei (detected with the *Smarcd3*-F6-nGFP reporter) during a collection of E7.25 – E7.5 time lapse series, using ventral partial maximum projections. Nuclei orientations were plotted as a function of their medial-lateral position in the crescent. **D**. Examination of JCF cell (arrowheads) molecular identities during MET of the cardiac crescent in a fixed E7.5 embryo in ventral max projection. **E**. UMAP shown in Fig. 5F, with superimposed labels corresponding to the stages as named in source publication [39]. **F**. The cluster markers are shown by heatmap. **G**. Notable markers are depicted in qualitative co-expression feature plots.

**Figure S6:**
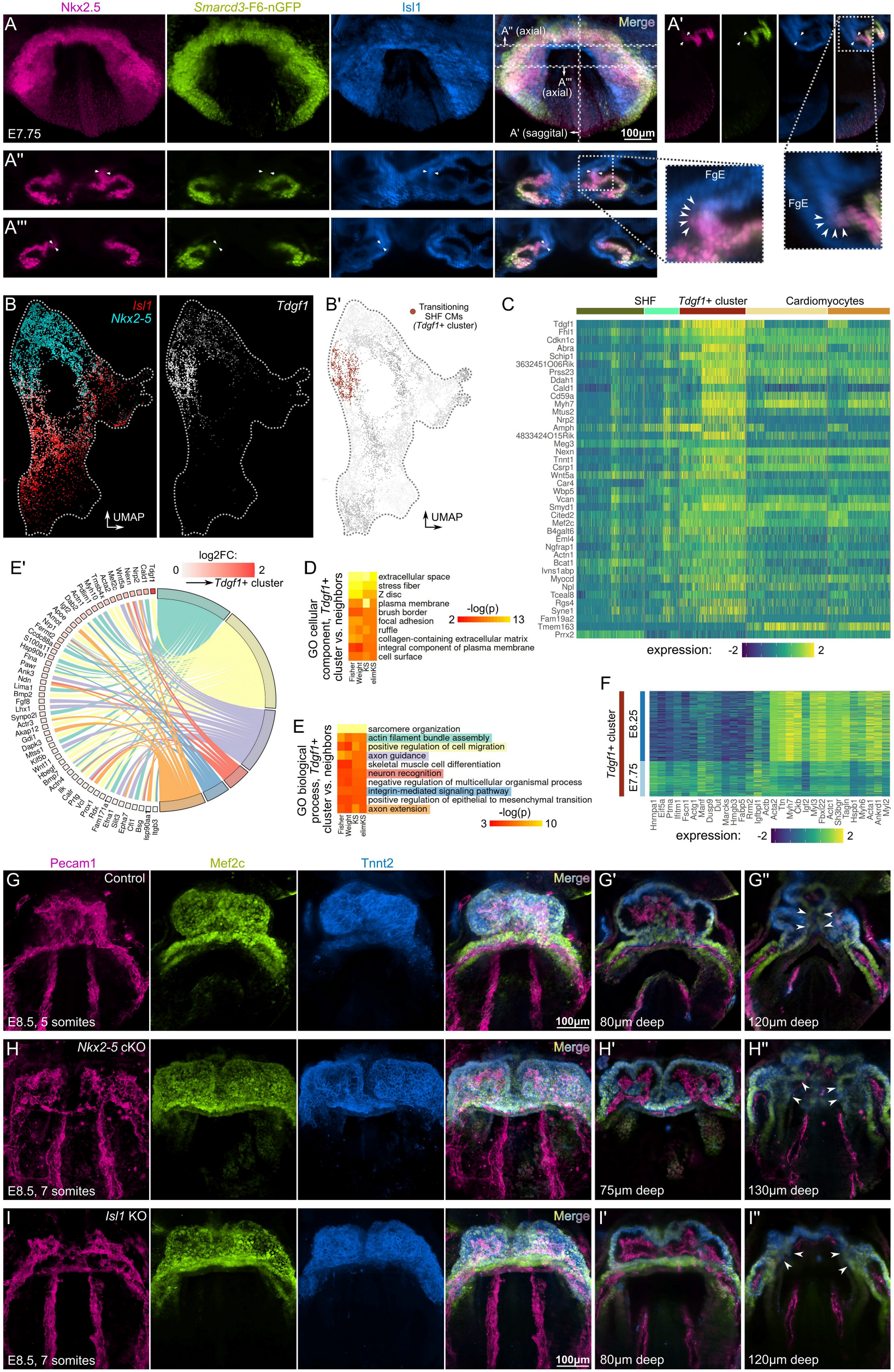
LHT closure by *Isl1*-dependent morphogenetic wave within differentiating SHF progenitors, related to Figure 6. **A-A”’**. LSFM examination of fixed embryos at E7.75, immediately prior to LHT formation. Max projection ventral views are shown (**A**). Midline sagittal plane (**A’**) and two axial planes (**A”** and **A”’**) are examined in 7.5μm thick slices. Arrowheads in **A-A”’** identify the morphogenetic wave within the boundary zone between ISL1 and NKX2.5 expression. **B-B”**. By scRNAseq [39], we identified the respective UMAP cluster (**B’**) by qualitative co-expression with *Nkx2-5*, *Isl1*, and *Tdgf1* (**B**). **C**. Differential expression of *Tdgf1*+ cluster and the four surrounding clusters, with lowest p-value differentially expressed genes plotted. **D-E’**. Positive-only (upregulated in *Tdgf1*+ cluster) significantly altered genes were assessed for gene ontology (GO) membership, with the most significant 10 categories shown for cellular component (CC, heatmap in **D**) or biological process (BP, heatmap in **E**). Six interesting BP terms are inspected by gene membership and log2FC differential expression (*Tdgf1*+ cluster versus four neighbors), as shown in chord plot (**E’**). **F**. The *Tdgf1*+ cluster is examined at E7.75 versus E8.25, with top log2FC positive and negative differentially expressed genes plotted. **G**-**I”**. Morphologic defects in LHT closure of *Isl1* KO and *Nkx2-5* cKO mice, at indicated stages for comparison. Ventral max projections are show in **G**, **H**, **I**, with cutaway 7.5μm thick ventral view slices show at indicated depths in **G’**, **G”**, **H’**, **H”**, **I’**, **I”**. Arrowheads (**G”**, **H”**, **I”**) point to normal dorsal seam / prospective dorsal mesocardium in control embryos (**G”**), with delayed dorsal closure in *Nkx2-5* cKO embryos (**H”**), and total absence of morphogenetic wave formation in *Isl1* KO embryos (**I”**).

**Figure S7:**
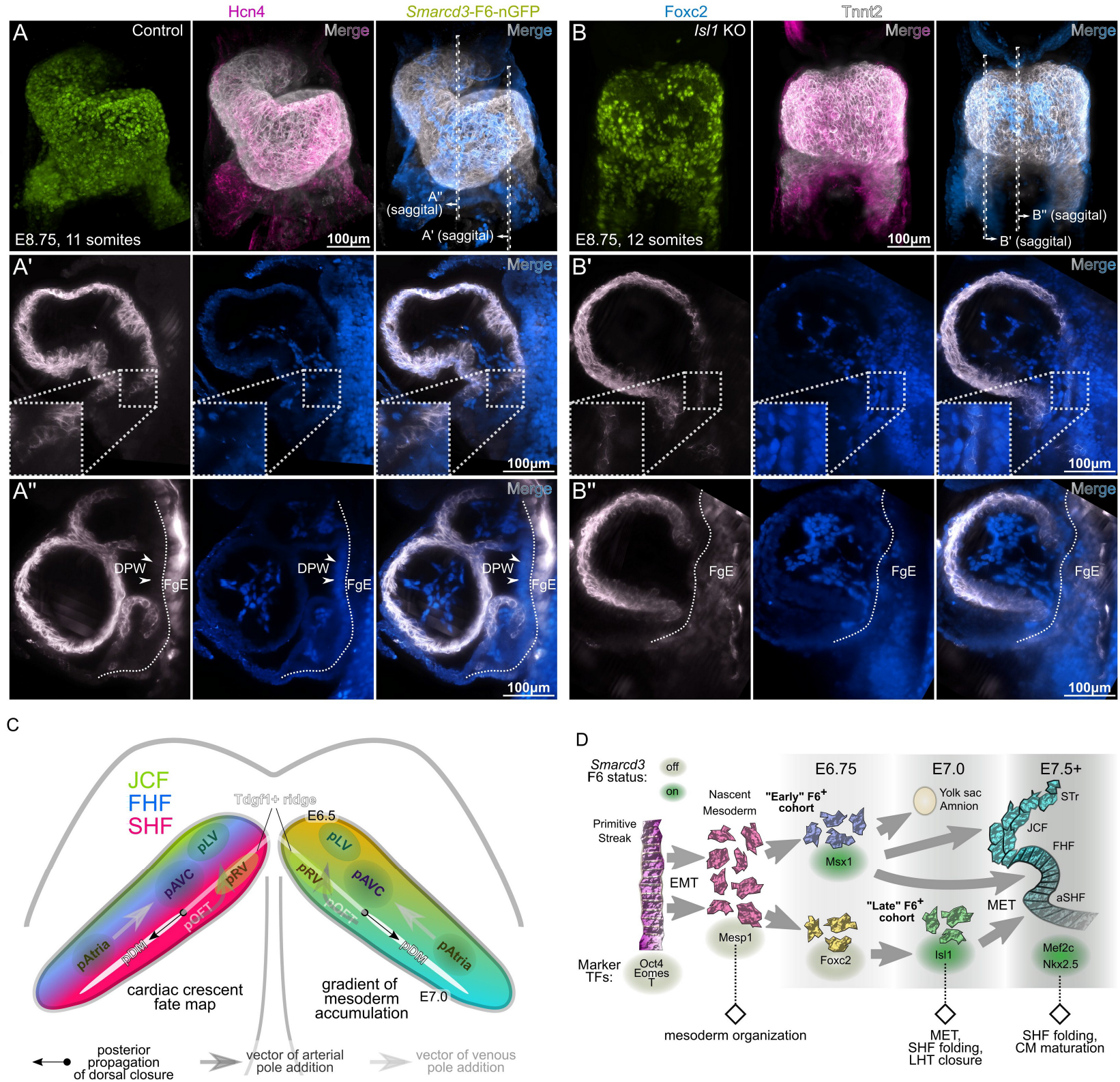
*Isl1*-dependent LHT closure and diagonal spatiotemporal gradient of gastrulation, related to Figures 6 and 7. **A-B”**. Control (**A-A”**), or *Isl1* KO (**B-B”**) embryos are examined for morphology by LSFM. Ventral view max projections are shown (**A** and **B**), as are 7.5μm thick sagittal cutaways (**A’**, **A”,B’**, **B”**) at locations indicated in A and B. Arrowheads point to dorsal pericardial wall (DPW) in control embryos (**A’**, **A”**), which is missing due to failed LHT closure in *Isl1* KO embryos (**B’**, **B”**). FgE = foregut endoderm. **C**. Spatiotemporal map of early cardiac specification, ventral schematic showing cardiac crescents. Left crescent depicts early cardiac fields, orientation longitudinally along anterior-posterior axis along crescent. Right crescent shows gradient of mesoderm accumulation, with progenitor birthdates (E6.5-E7.0) labeled at extremes. The *Tdgf1*+ ridge of transitioning SHF progenitors is shown (gray line), which forms the entry point for arterial pole addition. Black dot and arrow represent first point of closure of dorsal mesocardium, an event that propagates posteriorly in crescent (white line). pLV = prospective LV, pRV = prospective RV, pAVC = prospective AV canal, pAtria = prospective atria, pOFT = prospective outflow tract, pDM = prospective dorsal mesocardium. **D**. General timeline of cardiac specification by *Smarcd3*-F6 progenitors. Following EMT during early gastrulation, nascent cells belong to two cohorts as demarcated by MSX1 and FOXC2 labeling. The early F6^+^ cohort, being already present by E6.75, is mostly made up of MSX1^+^ cells, while the late F6^+^ cohort is probably a mixture of MSX1^+^ and FOXC2^+^ nascent progenitors, favoring the latter. The FHF and SHF undergo MET, forming the epithelium that folds into the early LHT.

## Materials and methods

### Study design and method details

#### Animal Subjects

All mouse protocols were approved by the Institutional Animal Care and Use Committee at UCSF. Mice were housed in a barrier animal facility with standard (12-hour dark/light) husbandry conditions at the Gladstone Institutes. *Smarcd3*-F6-nGFP and *Smarcd3*-F6-CreERT2 mice were described previously [6]. *Mesp1*-Cre knock-in mice [8,43] were obtained from Yumiko Saga. Cre/lox reporter lines RCL-H2B-mCherry and RCL-tdTomato (Ai14) are available at Jackson Laboratory (#023139 and #007914). *Mef2c*AHF-Cre mice were obtained from Brian Black [52]. *Isl1*-Cre and *Nkx2-5*-flox mice are available at Jackson Laboratory (#024242 and #030554). Mice for knockout experiments were maintained on a mixed CD-1 / C57BL/6J background, while control embryos for the majority of live imaging were generated by mating C57BL/6J males to CD-1 females. When indicated in figure panels, multiple reporter and/or mutant alleles may be present in the same embryo(s), either in isolation or in combinations of the following. *Smarcd3*-F6-nGFP refers to *Hipp11^Smarcd3^*^-F6-Hsp68-nGFP/+^. “*Smarcd3*-F6 lineage” denotes embryos with *Hipp11^Smarcd3^*^-F6-Hsp68-CreERT2/+^;*Rosa26*^CAG-LSL-tdTomato/+^. “*Mesp1* lineage” _denotes embryos with *Mesp1*_Cre/+_;*Rosa26*_CAG-LSL-H2BmCherry/+ _or with *Mesp1*_Cre/+_;*Rosa26*_CAG-LSL-tdTomato/+ genotypes. “*Mef2c*AHF lineage” denotes embryos with *Mef2c*AHF-Cre;*Rosa26*^CAG-LSL-tdTomato/+^ genotype. “*Mesp1* KO” or “*Mesp1*^-/-^” are embryos with *Mesp1*^Cre/Cre^;*Rosa26*^CAG-LSL-H2BmCherry/+^ genotype. “*Isl1* KO” denotes embryos with *Isl1*^Cre/Cre^;*Rosa26*^CAG-LSL-tdTomato/+^ genotype, for which matched controls are either *Isl1*^Cre/+^;*Rosa26*^CAG-LSL-tdTomato/+^ (where *Isl1* lineage is quantified), or *Isl1*^+/+^ (where *Isl1* KO is compared with other mutants). *Nkx2-5* cKO refers to embryos with *Mesp1*^Cre/+^;*Nkx2-5*^flox/flox^ genotype, for which matched controls are *Mesp1*^Cre/+^;*Nkx2-5*^+/+^. Following set up of timed matings, the day of copulatory plug is designated as E0.5. For each embryological process we wished to study, we imaged 3-4 embryos by time lapse LSFM, though we chose only those with the highest imaging quality for further analysis. In this work, we thus present only phenomena that were clearly reproduced over many embryos, on both subjective and quantitative bases.

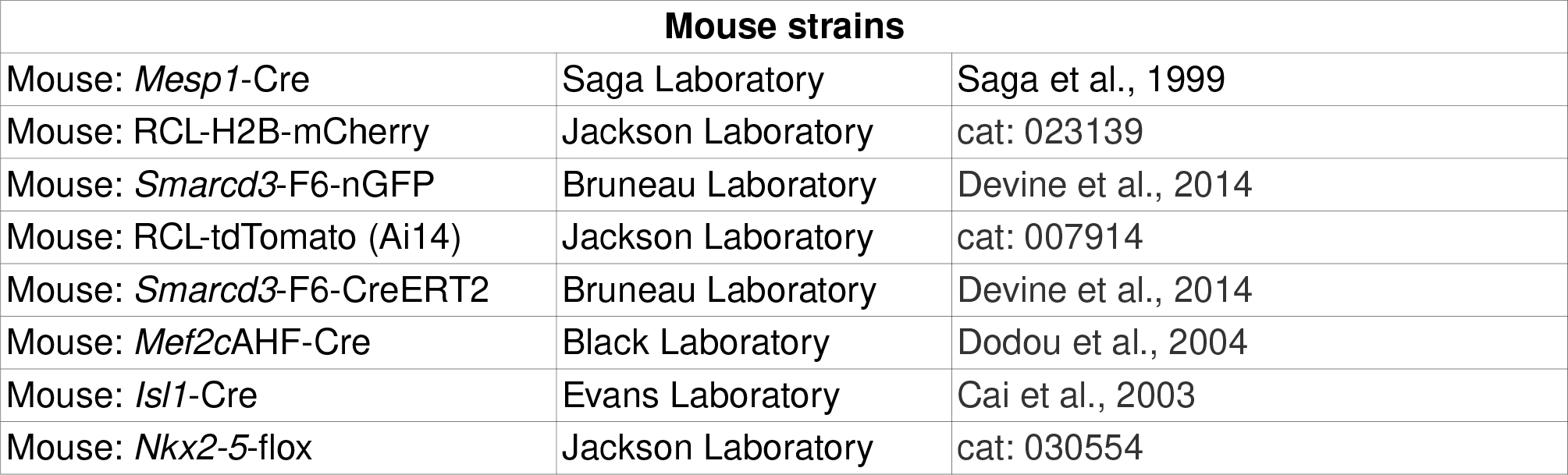

| Mouse strains |  |  |
| --- | --- | --- |
| Mouse: *Mesp1*-Cre | Saga Laboratory | Saga et al., 1999 |
| Mouse: RCL-H2B-mCherry | Jackson Laboratory | cat: 023139 |
| Mouse: *Smarcd3*-F6-nGFP | Bruneau Laboratory | Devine et al., 2014 |
| Mouse: RCL-tdTomato (Ai14) | Jackson Laboratory | cat: 007914 |
| Mouse: *Smarcd3*-F6-CreERT2 | Bruneau Laboratory | Devine et al., 2014 |
| Mouse: *Mef2c*AHF-Cre | Black Laboratory | Dodou et al., 2004 |
| Mouse: *Isl1*-Cre | Evans Laboratory | Cai et al., 2003 |
| Mouse: *Nkx2-5*-flox | Jackson Laboratory | cat: 030554 |

#### Whole embryo dissection and culture conditions

Pregnant dams were sacrificed on the day of the experiment, per institutional IACUC standard procedure, and were immediately dissected, with uterus transferred to warm DMEM/F-12 with HEPES and without phenol red. Gestational sacs were transferred to 37°C dissection medium (DMEM/F-12 w/ HEPES and w/o phenol red, 10% heat-inactivated fetal bovine serum, 1X penicillin-streptomycin, 1X ITS-X, 1X GlutaMAX, as well as 8 nM β-estradiol, 200 ng/ml progesterone, 25 μM N-acetyl-L-cysteine as per [53]) in small batches (4-5 per 6cm round bottom dish). While maintaining 37°C as best as possible, embryos were microdissected using fine forceps, and were transferred to 37°C culture medium (identical to dissection medium except with 50% of DMEM/F12 volume replaced by heat-inactivated rat serum, resulting in final 42.5% rat serum) using low-retention wide orifice pipette tips. Embryos were screened for reporter expression and morphology using a standard fluorescence dissection microscope (Leica). Embryo stage was determined with standardized methods [6], including the use of mouse embryo atlases, in combination with operator judgement for finely granular assessments.

#### Embryo preparation for live LSFM

Embryos were maintained in culture medium at 37°C and 5% CO_2_ until live imaging began. At the time of imaging for embryos at E7.5 and beyond, culture medium was supplemented with 2μM CB-DMB to decrease (but not obliterate) motion artifact from beating due to its activity on Ncx1 channels [54], for which genetic loss results in normal development with structurally normal hearts until at least E8.5 [55,56]. Before mounting, glass capillaries were pre-filled with liquid embedding medium (1.5% agarose, 3% gelatin in PBS, microwaved and mixed until fully melted) and pistons were inserted, then allowed to cool to ∼35°C before use. Using a stereoscopic dissection microscope (Leica), each free end (opposite the piston rod) of the embedding mix was extruded and 25-30% of its was length trimmed with a dissection forceps. Each embryo was attached by pushing its ectoplacental cone into the partially-gelled column. After confirming good attachment, the embryo and a small volume of surrounding culture medium were drawn inside the capillary and parked about 4-5mm from the open end. Capillaries were maintained at 37°C as best as possible until imaging.

| Reagents for live imaging |  |  |
| --- | --- | --- |
| Low MP agarose | Fisher | BP165-25 |
| Gelatin | Sigma | G1890 |
| Rat Serum, special collection | Valley Biomedical | AS3061-SC |
| Fetal bovine serum | ThermoFisher | 10082139 |
| DMEM/F-12 | ThermoFisher | 11039021 |
| GlutaMAX | ThermoFisher | 35050061 |
| ITS-X | ThermoFisher | 51500056 |
| Penicillin/Streptomycin | ThermoFisher | 15070063 |
| b-estradiol | Sigma | E8875 |
| Progesterone | Sigma | P3972 |
| N-acetyl cysteine | Sigma | A7250 |
| CB-DMB | Sigma | C5374 |
| Glass capillary and piston, largest | Sigma | Z328510 and BR701934 |
| Glass capillary and piston, large | Sigma | Z328502 and BR701938 |
| Glass capillary and piston, small | Sigma | Z328480 and BR701932 |
| Glass capillary and piston, smallest | Sigma | Z328472 and BR701930 |
| Wide orifice low-retention tips | Rainin | 30389197 |

#### Live LSFM imaging (Fig. 1A)

Lightsheet Z.1 (Zeiss) with incubation and dual pco.edge 4.2 cameras (PCO) was configured prior to embryo harvest, using a 20X/1.0 plan apochromat water-dipping detection objective with refractive index correction collar set to n=1.38, dual 10X/0.2 illumination objectives, and tank pre-filled with culture medium as described above at 37°C and 5% CO_2_. Embryo capillaries were auditioned for imaging quality and position, and chosen embryos subjected to 9-24 hours of LSFM imaging using our Zeiss Lightsheet Adaptive Position System (ZLAPS), linked in the key reagent table. ZLAPS is a user-friendly AutoIT GUI application that interfaces with ZEN, using multiview acquisition settings established by the user. We typically used 2-3 frontal views with 72° - 110° offsets, and collected GFP/488nm/505-545nm and RFP/561nm/570+ nm channels simultaneously. ZLAPS captures new images at fixed time intervals (specified by the user), and calls ImageJ with the Java SIFT [57] plugin to register sequential acquisitions. The registration matrix outputted by SIFT (for each view) is used to adjust (with hysteresis and over-correction mitigation) the stage position of the Z.1 for subsequent acquisitions. For long-term imaging (24hr+), additional optimizations are necessary: light sheet alignment is checked and manually adjusted every 4-6 hours, piston rods are secured with Parafilm, the specimen tank/chamber cover is used, and additional sterile water and/or culture medium is trickled/dripped (< 0.5mL/hour) into the tank using a micro-osmotic pump to overcome evaporation losses.

#### Whole mount preparation for fixed LSFM imaging

Embryos were harvested as for live imaging, except uterus was transported and dissected in ice cold PBS. Embryos were fixed in 4% paraformaldehyde for 1 hour at room temperature with gentle agitation, and washed briefly in PBS before being transferred to short-term storage at 4°C in PBS with 0.2% sodium azide. For immunostaining, embryos were transferred individually to wells of PCR tube strips. E9.5 embryos were cleared briefly in 8% SDS in 200mM borate buffer [58], with gentle agitation for a few hours at 37°C until clear, followed by 2-3 washes in PBS at 37°C. Smaller embryos were not subjected to clearing. Subsequently, embryos were incubated in blocking solution (PBS with 5% normal donkey serum, 0.2% sodium azide, and 0.5% TX-100 for E5-E7 embryos, 0.65% for E7-E8, 0.8% for E9) plus 100μg/mL of unconjugated Fab fragment donkey anti-mouse, for 2 hours at 37°C with gentle rocking/rotation. After washing in PBS, primary staining was performed in blocking solution overnight, followed by additional washing. Secondary incubation was performed in blocking solution for 2-3 hours, followed by final washing, with all steps at 37°C with gentle rocking/rotation. For storage at 4°C until mounting, labeled E6-E7 embryos were sunk in 40% glycerol in PBS, while older embryos were kept in PBS.

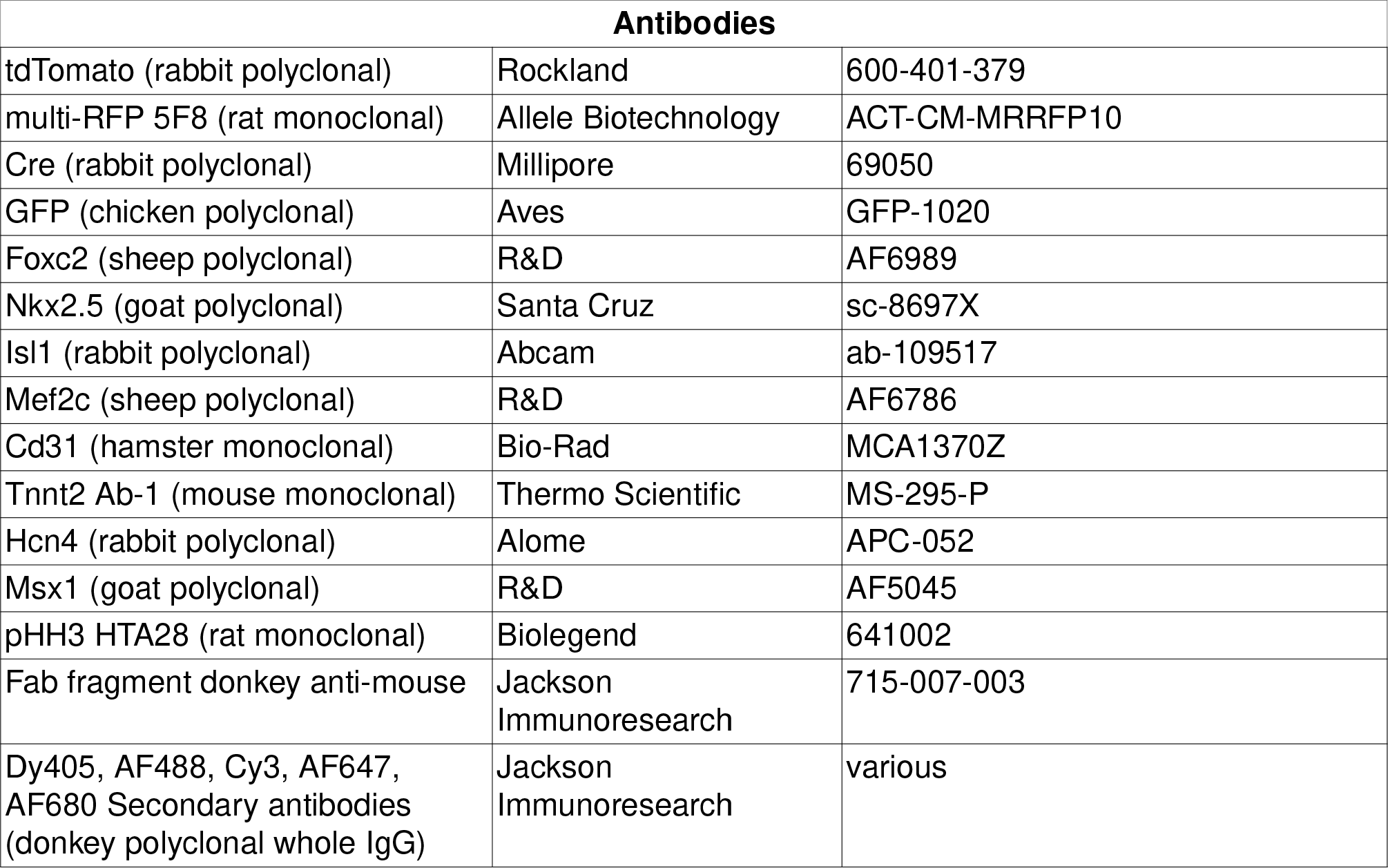

#### Fixed LSFM imaging

Embedding medium (2% agarose in PBS) was melted in a microwave and cooled to 35°C, when embryo(s) were immersed for 30 seconds with gentle mixing. Glass capillaries were partially filled with liquid embedding medium, and their pistons were retracted to pick up embryos. Following cooling and gelling of the embedded embryos, capillaries were taped to the inside walls of polystyrene tubes, and specimens were extended into room-temperature immersion medium (EasyIndex OCS for E8+ embryos, or 40% glycerol for E6-E7 embryos) for overnight equilibration. Specimens were imaged on Lightsheet Z.1 (Zeiss) with dual pco.edge 4.2 cameras (PCO) for simultaneous two-channel acquisition using standard illumination lasers (405nm, 488nm, 561nm, 638nm). Rarely, channel bleed necessitated later subtraction during processing. Three views were acquired from the ventral aspect of each specimen at 72° (E7.5+) or 90° (E6.5-E7.25) offsets, using 20X/1.0 plan apochromat water-dipping detection objective at n=1.38 for 40% glycerol immersion (mated with 10X/0.2 “LSFM” illumination objectives), or 20X/1.0 plan neofluar clearing dipping objective at n=1.45 for EasyIndex OCS immersion (mated with 10X/0.2 “LSFM clearing” illumination objectives).

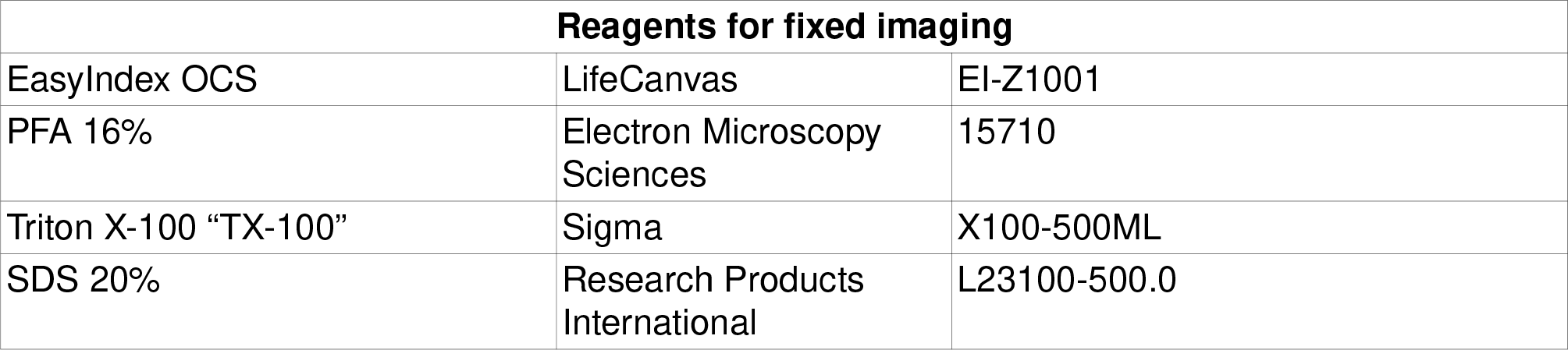

| Reagents for fixed imaging |  |  |
| --- | --- | --- |
| EasyIndex OCS | LifeCanvas | EI-Z1001 |
| PFA 16% | Electron Microscopy Sciences | 15710 |
| Triton X-100 “TX-100” | Sigma | X100-500ML |
| SDS 20% | Research Products International | L23100-500.0 |

#### Computer hardware and software environment

ZEN and Lightsheet Z.1 acquisitions were run on a Zeiss-supplied workstation with dual 8-core 2^nd^ generation Intel Core based Xeon processors and 96GB RAM, running Windows 7. Data was processed on workstations with either single 8-core 10^th^ generation or dual 8-core 3^rd^ generation Intel Core based Xeon CPUs, 128GB RAM, and 4GB Nvidia GTX 1650 GPUs, running Kubuntu 20.04 LTS with Nvidia driver 470, Fiji v2.1.1, Python 3.8.10, Perl 5.30.0, and CUDA toolkit 11.1. All software-comparative benchmarks were run on the same system. Accuracy evaluations between TGMM versions were performed by running each version with its optimized parameter set (determined empirically through iterative comparison), followed by import to MaMuT. Random subsets of cells, tracks, and divisions were assessed in single and double-blinded fashion, with annotations made and counted using MaMuT Perl scripts. Single cell RNAseq analysis was performed on similar hardware running Kubuntu, RStudio desktop build 443, r-base 4.1.3, Seurat 4.0.6 [59], topGO 2.48.0, and GOplot 1.02.2 [60].

#### Raw image processing and single view deconvolution (Fig. 1B)

ZEN-generated .czi files were handled with our CZI LSFM Processing Scripts (see Software table in Materials and methods) in Fiji [34]. The initial step (“deconvolve .czi files”) batch processes live or fixed raw data. First, a theoretical point spread function (PSF) is generated, based on illumination and detection parameters (as the intersection of Gaussian light sheet with modeled widefield detection) embedded in Zeiss metadata, with an optional detection NA penalty for the improved aberration handling. Each channel of each view is deconvolved for each timepoint, using a closed form solution with Tikhonov regularization [33]. We had determined this approach was the best balance of result quality and computational intensiveness, following extensive empirical testing and benchmarking with a wide range of fixed and live samples. After .tif files are written for each channel, view, and timepoint, additional automated filtering (“filter LSFM .tif files”) is performed that can include (by user preference) background subtraction deblurring, bright blob and/or precipitate removal, bit depth compression, z-stack depth equalization (needed for BigStitcher), and/or maximal intensity projection export. Because the many serially-performed functions have user-controllable settings, changes or alterations to the output images may be somewhat unpredictable or unnatural. For new experiments, we recommend a trial-and-error approach to determine the best protocol. We typically handled fixed image datasets at 16-bit depth with maximal automated filtering including bright blob removal (helpful for deep max Z projections in whole mount IHC), although frequent artifacts remain. Live datasets, on the other hand, were usually contrast-enhanced uniformly across each entire 4d stack, then range-compressed to 8-bit.

#### Multiview alignment and fusion (Fig. 1C)

After deconvolution and filtering, resultant .tif files were imported into BigStitcher [28], using its automatic loader. “Interest points” were detected within one or more channels, across all views and timepoints, and views were registered in 3d followed by 4d space. The most optimal solution for live datasets resulted from pre-registration using a “Fast Descriptor-Based” method in 3d then 4d, followed by drift mitigation in time with a “Center-of-mass” method, followed by “Fast Descriptor-Based” or “Precise descriptor-based” methods on the whole dataset and in regions of difficulty. Finally, multiple “Iterative closest point” steps were used to improve upon remaining view-to-view and timepoint-to-timepoint offsets. Multiview fusion was performed using optimized “lightweight” content-based fusion, coded within our fork of BigStitcher’s multiview-registration plugin (see Software table within Materials and methods). Other advantages of our forked plugin include fusion in multiple axes, and use of an arbitrary z-anisotropy factor (we use 4). Following fusion into single image volumes, datasets can be viewed in BigDataViewer in Fiji, or can be further processed in batch using additional components of our CZI LSFM Processing Scripts. This includes automated generation of oblique 3d projections, as well as single-channel anaglyphs (Video S7).

#### F-TGMM v2.5

Tracking with Gaussian Mixture Models (TGMM) 1.0 [61], and its successor TGMM 2.0 [29], are open-source packages for analysis of large-scale time-lapse cellular imaging. With linear best-fit modeling (from one timepoint to the next) of a whole-specimen Gaussian mixture, TGMM is fast and accurate. It is written in C++, and utilizes GPU/GPGPU acceleration in CUDA to perform several critical steps. TGMM’s accuracy owes itself to several factors: 1. watershed hierarchical segmentation for identifying 3d supervoxels (i.e. Gaussians / prospective cells) – which is superior to a difference-of-gaussians approach as in Trackmate [62]; and 2. the implementation of “temporal logical rules,” which build on the linear model by extending false cell deaths, and connecting new births to prospective division parents. We modified TGMM to enhance its performance of with our data. First, over- and under-segmentation were improved by applying dynamic rather than static “background subtraction” to the input images, using Gaussian-blurring (user configurable) to define background. Second, we liberalized the dead cell extension rules to further improve linkage across time. Third, we re-wrote the cell division classifier, which was constrained to calling ‘yes’ or ‘no’ on division trios already assigned by the linear model. Instead, our new classifier incrementally improves division linkage accuracy by sampling trios in the neighborhood of each new birth, and assigning scores to each one. Fourth, we re-wrote the main tracking loop to eliminate repeat calls to hierarchical segmentation for the same image, instead caching the result within the temporal window (usually ±5 timepoints) for re-use. Last, we fixed a number of bugs, streamlined the code’s output to stdout, and made updates necessary for compiling and running on contemporary CUDA hardware and software. Overall, a complete TGMM v2.5 run is typically 30% faster than TGMM 2.0, and produces more accurate results. Regrettably, division classification is still suboptimal even with the above improvements and iterative training of the classifier. Notably, we could not run the convolutional neural network (CNN) division detector included with TGMM 2.0 [29] outside of its Docker container, and even there it produced extremely poor results with our datasets. Much work remains in the arena of automated division detection, including not just the identification of division events, but in linking the correct daughter pair to each mother.

#### Tracking at single cell resolutions (Fig. 1D)

Fused image volumes of *Mesp1* lineage, from either the front or side view of each embryo, were used as input for tracking. A python script “bdv_export_all_h5_to_klb_pyklb.py,” included with F-TGMM v2.5, converts the fused output from .h5 format to .klb format [29], making it compatible for input with both BigDataViewer and F-TGMM. The empirically-determined optimal F-TGMM configuration parameters used on our datasets are provided in the below table. ProcessStack was run individually (scripted for batch processing) for watershed segmentation of each timepoint’s fused volume, followed by a single TGMM call on the entire dataset. Rare, sporadic, dropout of cell linkages were corrected on the resulting TGMM .xml data using a perl script “XMLfinalResult_fix_cell_NaNs.pl.”

| Parameter | TGMM 2.0 | F-TGMM v2.5 |
| --- | --- | --- |
| backgroundThreshold | 5 | 5 |
| radiusMedianFilter | 2 | 2 |
| sigmaGaussianBlurBackground | N/A | 20 |
| useBlurredImageForBackgroundDetection | N/A | 0.7 |
| weightBlurredImageSubtract | N/A | 0.6 |
| minTau | 0 | 0 |
| persistanceSegmentationTau | 1 | 1 |
| betaPercentageOfN_k | 1.6 | 2.5 |
| nuPercentageOfN_k | 0.5 | 0.1 |
| alphaPercentage | 0.85 | 0.75 |
| maxIterEM | 50 | 50 |
| tolLikelihood | 1e-6 | 1e-6 |
| regularizePrecisionMatrixConstants_lambdaMin | 0.05 | 0.02 |
| regularizePrecisionMatrixConstants_lambdaMax | 0.8 | 0.8 |
| regularizePrecisionMatrixConstants_maxExcentricity | 16.0 | 16.0 |
| temporalWindowForLogicalRules | 5 | 5 |
| SLD_lengthTMthr | 5 | 5 |
| conn3D | 74 | 74 |
| minNucleiSize | 1600 | 1200 |
| maxNucleiSize | 20000 | 10000 |
| maxPercentileTrimSV | 0.55 | 0.8 |
| conn3DsvTrim | 6 | 6 |
| maxNumKNNsupervoxel | 10 | 10 |
| maxDistKNNsupervoxel | 40 | 40 |
| thrSplitScore | -1 | -1 |
| thrCellDivisionPlaneDistance | 14 | 14 |
| cellDivisionClassifierMethod | AmatF2013 | DominguezM2021 |
| thrCellDivisionWithTemporalWindow | 0.45 | 0.45 |

#### Mining and analysis of tracking data (Fig. 1E)

F-TGMM writes .xml tracking solutions, representing the linkages that connect each cell to its past and future self across time. These tracks can be imported directly into MaMuT [30], a Fiji plugin for annotation and visualization of big datasets. Our fork of the MaMuT plugin (see Software table in Materials and methods) contains fixes to the TGMM import code, enables track vector viewing in 2d, and makes a number of improvements in MaMuT’s 3d viewer for better performance with large datasets, although this feature was recently removed in the upstream mainline repository. Moreover, we have written a large compendium of scripts in Perl for filtering, labeling, subsetting, motion subtracting, merging, analyzing, and exporting from MaMuT datasets, the features of which are not available in the mainline plugin. These scripts were employed in various operational workflows, for generating the many viewable and analyzable MaMuT datasets presented in this work. Lastly, we updated the SVF package [29] with bug fixes and for use with Python3. Where indicated, we processed TGMM data with SVF to generate long-running vector fields of the dataset for morphometric assessment, which facilitated an overall understanding of tissue deformation during heart development. When individual tracks at single-cell resolution were desired (SVF not indicated), we typically filtered datasets for tracks of 2-4 hour minimum length, without abrupt unnatural movements, and occasionally would manually remove tracks not belonging to the cell type of interest. Starting with a MaMuT .xml dataset (derived either directly from TGMM or via SVF), included scripts facilitate export of spacetime coordinates for each track, which were summarized for statistical analysis in spreadsheet software or R.

#### Single cell RNAseq analysis

Single cell wild-type datasets [39,54] were downloaded from public repositories, and analyzed in Seurat [59] v4.0. We tailored the dataset normalization and integration method (CCA, SCtransform) to specific batch effects and coalescence of like clusters in UMAP space. Initial QC cleanup involved removal of low quality cells, and those belonging to either endoderm or ectoderm lineages. Subsequent clusters were subsetted to depict only pre-cardiac mesoderm and its derivatives. All differential gene expression analysis was performed with FindMarkers in Seurat, and lists of differentially-expressed and non-differentially-expressed genes were inputted into topGO for gene ontology analysis. Pearson correlation was performed on normalized RNA count data. Result visualization was scripted with ggplot2, Seurat, GOplot [60], and/or igraph. Qualitative co-expression feature plots were generated by overlay and assignment of individual feature plots to different channels in Fiji. A single cell *Mesp1* KO dataset was generated for a companion manuscript (Krup et al., manuscript in preparation), and was analyzed for differential expression of select features relevant to directional migration of mesoderm [45] and related signaling.

### Data analysis

#### Mesoderm accumulation

Using direct TGMM imported data and MaMuT scripts, we parsed each embryo (E6.5 – E7.0) into 9 bins comprised of a 3 × 3 rectangle box pattern as seen in the lateral view, and filtered for QC as described above. For quantification smoothing, each box shares an overlap of 50% of the nearest tracks in each adjacent neighboring box(es). Track birthdate is the timepoint of first appearance of the track. Track density, for each cell within a bin, is the number of other cells present within a radius spanning 12 times the radius of that cell. Track motility was computed as the average of all moving window velocities for a discrete time span (i.e. 30 minutes), incremented each frame over the life of that track. SVF analysis was performed for tissues (i.e. embryonic mesoderm and extraembryonic mesoderm) assigned and painted within SVF’s tissue-bw script. Track mean velocity is the total distance traveled divided by the total time span of the track, and is of particular use with SVF analyses. Mean comparisons were based on Welch t-test.

#### Assessments of cell neighbor relationships and mixing

For quantification of separation after cell division, an empty MaMuT dataset was manually annotated with division nodes and daughter tracks derived from a random assortment of such events in each fused BigDataViewer dataset. Using a custom Perl script to analyze the MaMuT datasets, mother and daughter positions were exported into a table. Raw measurements were also indexed to a singular length of an average embryo from this stage. For quantification of track position exchanges, we separated each embryo into two bins by cell proximal-distal position, then again by lateral half, resulting in four bins for analysis. We used a custom Perl script to analyze tracks in pairwise fashion within the MaMuT datasets, bounded by time and cell distance cutoffs as specified by user (here, co-existent tracks were admitted until t+4.5h into the dataset, and rejected if they were separated by more than 250μm distance in the axis of analysis). Each pair is assessed for its distance offset in the dimension of interest, and those distances can be compared over time to determine whether the tracks exchange position in that dimension. End offsets were first plotted as a function of begin offset, and the relationship was assessed by Pearson correlation coefficient R^2^. Next, the offsets were followed in time to determine the number of position exchanges along the axis, and the average number was plotted for each bin and axis. All mean comparisons as described above were made by Welch t-test.

#### Birth of Smarcd3-F6 progenitors

In order to bin tracking results by *Smarcd3*-F6 status, F-TGMM tracking solutions for *Mesp1* lineage progenitors at E7.0 were processed with SVF, using the *Smarcd3*-F6-nGFP channel as a mask for tissue-bw. When tissue-bw was performed for early (forward propagation) and late (backwards propagation) timepoints, different sets of tracks were included in the F6^+^ pool, though late tracks almost always included early tracks as a subset. Using MaMuT Perl scripts, we subtracted the early tracks from the late tracks, and colored all tracks by F6 status: off, on early, or on late (which included the vast majority of on early tracks). Complete painted solutions were visualized with MaMuT. They were also subjected to uniform sparsification (via Perl script) and plotted as orthographic projections to depict characteristic migration patterns.

#### Cell fates of the Smarcd3-F6 lineage

Lineage analysis was carried out in fixed embryos imaged by LSFM as described above. Using fused image volumes, we attempted to count all cells in all embryos, assigning them to myocardial or non-myocardial structures. Comparison of their mean contributions to various structures was made by Welch t-test.

#### Counting Mesp1 lineage and Smarcd3-F6 progenitors

Counts of *Mesp1* lineage progenitors were made using live LSFM datasets that had been tracked with F-TGMM, using the number of tracked cells at corresponding timepoints as the initial estimate. Those estimates were further refined by subtracting estimated incidentally-labeled cells (i.e. endoderm, etc). *Smarcd*-F6-nGFP counts were made by performing background subtraction in Fiji with kernel size 50, then by examining corresponding timepoints with Trackmate’s DoG detector with radius 15 and threshold 5. Density of the DoG detection solution was determined by counting number of cells within an arbitrary radius of each cell (i.e. 20μm).

#### Cell morphometry during cardiac crescent MET

The volume of *Smarcd3*-F6 progenitors was estimated using a custom ImageJ macro, which evaluated *Smarcd3*-F6-nGFP and whole-cell tdTomato (*Mesp1* lineage) for a number of timepoints. In brief, the macro performs dilate alterations and background thresholding on the nGFP channel to create “spheres of influence” around each cell, which are then used as an intersect mask with the tdTomato channel. The intersection is measured for integrated intensity, which is divided by the estimated number of cells to yield estimated cell volume. Thickness of the overall crescent was estimated with manual measurements taken in sagittal plane slices. Cell density is summated for each cell as the number of cell neighbors within a stated radius, which is then averaged at individual time lapse frames near stated timepoints.

#### Quantifying movement behavior of the heart fields

After cardiac crescent MET, tissues and their descendant structures are revealed morphologically, allowing for F-TGMM tracking solutions to be subsetted into those constituent tissues via SVF. The tracks’ beginnings were tracked in reverse (i.e. via backward propagation), allowing for an assessment of sites of origin of the three layers principal layers derived during MET (pericardial, myocardial, and endocardial). Myocardial and pro-epicardial fields were analyzed for net track displacement, which could be assayed with or without the application of correction for (i.e. subtraction of nearby) endoderm movement by MaMuT Perl script. Endoderm correction was especially helpful during foregut folding and involution. For JCF position and motility assessments, we manually quantified F6^+^ cells in maximal z projections, because SVF agglomerates movements into vector fields, destroying nonuniform motility. Nuclei orientations were compared using Watson U_2_ test, whereas all other measurements were compared as means by Welch t-test.

#### Comparing Mesp1 mutants with controls

Since we had already determined that mesoderm accumulation occurs by diagonal spatiotemporal gradient, we compared cells from mutant and control embryos by assigning them to a position along that axis (rather than by membership to 3 × 3 spatial grids). We applied similar metrics utilized previously. Motility was computed as the average displacement sampled over 30-minute moving windows, and cell density as the average number of other cells counted within a 12-nuclei-radius from each cell. Additionally, track trajectories were scored in the lateral view, using the start and end coordinates to determine the directionality (in the orthographic lateral view). Trajectory angles (anterior at 0°, proximal at 90°) were calculated for each track in the orthographic lateral view, using a 2d vector from its start coordinate to end coordinate. Density distributions of all trajectory angles were plotted in polar space, and were compared with Watson U_2_ tests. All other measurements were compared as means by Welch t-test.

### Data and software

All software utilized to handle images, generate and process tracking solutions, and export data tables for analysis with R are available on Github, as listed below. Source data tables and R scripts used to generate individual figure panels are freely available from the authors upon request.

| Software |  |  |
| --- | --- | --- |
| Seurat 4.0 (R 4.0) | Satija Lab | Hao et al., 2021 |
| ggplot2 (R 4.0) | RStudio | Wickham, 2016 |
| TopGO 2.48.0 (R 4.0) | Alexa and Rahnenfuher | Alexa and Rahnenfuher, 2022 |
| GOplot 1.0.2 (R 4.0) | Ricote Lab | Walter et al., 2015 |
| F-TGMM v2.5 | This paper (e) and Fernando Amat | https://github.com/mhdominguez/F-TGMM |
| ZLAPS (ZEN lightsheet adaptive positioning system) | This paper (a) | https://github.com/mhdominguez/ZLAPS |
| TGMM2SVF | This paper (g) and Leo Guignard | https://github.com/mhdominguez/SVF |
| SVF2MaMuT | This paper (g) and Leo Guignard | https://github.com/mhdominguez/SVF2MaMuT |
| Fiji (base ImageJ v1.53f) | Schindelin, Rueden, Rasband, et al. | Schindelin et al., 2012 |
| - PSF Generator | Biomedical Imaging Group at EPFL | http://bigwww.epfl.ch/algorithms/psfgenerator/ |
| - Parallel Spectral Deconvolution | Piotr Wendykier | https://sites.google.com/site/piotrwendykier/software/deconvolution/parallelspectraldeconvolution |
| - CZI LSFM Processing Scripts | This paper (b and d) | https://github.com/mhdominguez/LSFMProcessing |
| - BigStitcher | This paper (c) and Preibisch Lab | Fiji update repositories and https://github.com/mhdominguez/multi-view-reconstruction |
| - KLB file format | Keller Lab | McDole et al., 2018 and Fiji update repositories |
| - MaMuT | This paper (f) and Tinevez, Pietzsch, et al. | Fiji update repositories and https://github.com/mhdominguez/MaMuT |
| MaMuT script library | This paper (h) | https://github.com/mhdominguez/MaMuTLibrary |

## Supplemental Video Titles

Video S1: Spatiotemporal assembly of mesoderm, related to Figure 2

Video S2: Birth of the Smarcd3-F6 cardiac progenitors, related to Figure 3

Video S3: Mesenchymal-epithelial transition of the cardiac crescent, related to Figure 4

Video S4: JCF motility and heart field morphogenesis, related to Figure 5

Video S5: Early heart tube formation, related to Figure 6

Video S6: Mesoderm assembly in *Mesp1* mutants, related to Figure 7

Video S7: Anaglyph 3d movies of early cardiac development, related to Figure S7

